# Emergent evolutionary forces in spatial models of microbial growth in the human gut microbiota

**DOI:** 10.1101/2021.07.15.452569

**Authors:** Olivia M. Ghosh, Benjamin H. Good

**Affiliations:** Department of Physics, Stanford University, Stanford, CA 94305; Department of Applied Physics, Stanford University, Stanford, CA 94305

## Abstract

The genetic composition of the gut microbiota is constantly reshaped by ecological and evolutionary forces. These strain-level dynamics can be challenging to understand because they emerge from complex spatial growth processes that take place within a host. Here we introduce a general population genetic framework to predict how stochastic evolutionary forces emerge from simple models of microbial growth in spatially extended environments like the intestinal lumen. Our framework shows how fluid flow and longitudinal variation in growth rate combine to shape the frequencies of genetic variants in sequenced fecal samples, yielding analytical expressions for the effective generation times, selection coefficients, and rates of genetic drift. We find that the emergent evolutionary dynamics can often be captured by well-mixed models that lack explicit spatial structure, even when there is substantial spatial variation in species-level composition. By applying these results to the human colon, we find that continuous fluid flow and simple forms of wall growth are unlikely to create sufficient bottlenecks to allow large fluctuations in mutant frequencies within a host. We also find that the effective generation times may be significantly shorter than expected from traditional average growth rate estimates. Our results provide a starting point for quantifying genetic turnover in spatially extended settings like the gut microbiota, and may be relevant for other microbial ecosystems where unidirectional fluid flow plays an important role.

---

The human gut harbors a diverse microbial ecosystem that plays an important role in maintaining health and reducing susceptibility to disease (1, 2). The composition of the gut microbiota is shaped by interactions between hundreds of different species (3, 4), which must continually reproduce in order to balance their daily excretion in feces (5, 6). These high rates of cell turnover create enormous opportunities for rapid evolutionary change: the trillions of bacteria within a single human colon produce more than a billion new mutations every day (7), and tens of thousands of bacterial generations will typically elapse within a single human lifetime (1, 8). Consistent with these estimates, a growing number of empirical studies, ranging from experimental evolution in mouse models (9, 10, 11, 12) to longitudinal sequencing of human subjects (7, 13, 14, 15, 16, 17, 18, 19, 20), have shown that genetic variants can sweep through resident populations of gut bacteria on timescales of weeks and months. Understanding the causes and consequences of this genetic turnover is likely to be important for predicting the stability and resilience of this diverse microbial ecosystem (21).

A central challenge in understanding the evolution of the gut microbiota stems from the complex structure of the gut environment. While the frequencies of genetic variants are typically tracked in homogenized fecal samples, most microbial growth occurs in the spatially extended setting of the large intestine. At present, little is known about how this spatial structure influences the evolutionary dynamics that are observed in sequenced fecal samples. This makes it difficult to draw quantitative conclusions from existing genetic data: What does it mean if the frequency of a variant increases dramatically over a few days? or if it eventually declines in frequency a few weeks later? Is it reasonable to interpret these coarse-grained measurements using well-mixed models from population genetics? or are spatially explicit models necessary for extracting quantitative insights?

Previous studies of gut microbial biogeography have shown that growth conditions can vary over a huge range of length scales, ranging from the micron scales of intercellular interactions (22, 23, 24) to macroscopic compartments like the mucosa, lumen, and intestinal crypts (25). Even within the comparatively simpler environment of the central lumen — thought to be the primary site of growth for some of the most abundant species of *Bacteroides* and *Firmicutes* (26) — significant longitudinal gradients are established by the basic anatomy of the intestinal tract. Primary nutrients are supplied at one end of the colon, where microbial densities are relatively low, and are rapidly consumed by bacteria in the ascending colon (27). The constant influx of fluid, along with additional mixing from contractions of the intestinal walls (26, 28), causes both bacteria and their metabolic byproducts to drift toward lower regions of the colon, where they are eventually excreted in feces. These physical processes combine to produce large gradients in nutrient concentrations, pH, and excretion rates across the longitudinal axis of the large intestine (27, 28, 29), which can have important consequences for the relative growth rates and abundances of different species (28, 30, 31, 32, 33, 34).

The evolutionary implications of these spatial gradients are less well understood, since the details will depend on the phenotypic differences that exist among more closely related strains. However, even in the absence of phenotypic differences, the longitudinal structure of the lumen will lead to general physical effects that can be appreciated by revisiting our back-of-the-envelope calculation for the production rate of new mutations. While billions of mutations may be produced each day, not all of these emerge on equal footing. Mutations that occur near the lower end of the colon are very likely to be excreted in feces before they can produce viable offspring. Conversely, mutations that occur near the entrance of the colon are significantly more likely to produce descendants that can persist and grow within the host over many generations.

These physical considerations suggest that only a subset of the luminal population will tend to contribute to its long-term genetic composition. Similar arguments apply for estimates of the effective generation time, which can vary substantially in different regions of the colon (27, 35, 36). These arguments suggest that the relevant evolutionary parameters must emerge from some sort of average over this existing longitudinal structure. However, the weights in this average will be determined not by the current bacterial densities and growth rates, but rather by their long-term contribution to the future population. Our limited understanding of these weights — even in the simplest models of luminal growth — leaves many important questions unresolved: Can fluid flow alone generate sufficient population bottlenecks that would allow large fluctuations in mutant frequencies within a host? How do absolute growth rate differences map on to the frequency trajectories that can be observed in vivo? And how do the answers to these questions depend on the physical properties of the gut environment, which vary dramatically across species, and can be modulated by drugs or other dietary perturbations?

In this study, we introduce a mathematical framework for predicting how stochastic evolutionary forces emerge from explicit spatial models of microbial growth in the intestinal lumen. Building on earlier work in Refs. (37, 38, 39, 40), we show that the emergent evolutionary dynamics can often be captured by spatially homogeneous models with an appropriate set of effective parameters, even when the ecological effects of spatial structure are very strong. We derive analytical expressions to show how the effective selection coefficients, generation times, and population sizes are related to the underlying growth rates and flow profiles along the large intestine. By applying this framework to parameter estimates from the human colon (28), we find that fluid flow alone is unlikely to generate large fluctuations in the frequencies of sequenced mutations, while the diffusion of slowly growing cells can still have a large impact on the long-term evolution of the population. These results provide a framework for understanding how spatial structure influences the genetic composition of the gut microbiota, and may be relevant for other microbial ecosystems (e.g. hot springs and acid mine drainage systems) where unidirectional flow is thought to play an important role (41, 42).

## ANALYSIS

We consider the dynamics of mutant lineages in a spatial model of luminal growth that incorporates continuous fluid flow, growth, and active mixing from contractions of the intestinal walls (26, 28). A key assumption of this model is that the mixing process is strong enough to destroy any cross-sectional spatial structure within the lumen (28), yielding an effective diffusion process in the longitudinal dimension (denoted by *x*) with some effective flow rate *v*(*x*) and effective diffusion coefficient *D*(*x*) (Fig. 1A). The local density of species *μ* at position *x* and time *t* can then be described by a reaction diffusion equation,

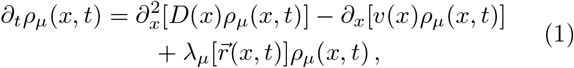

where 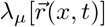 is a function that describes the net growth rate of species *μ*. We assume that this net growth rate depends on a set of local environmental variables 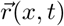, which could represent local nutrient concentrations, pH, or the abundances of other species. These environmental variables will be described by their own set of reaction diffusion equations, which are coupled to the species densities *ρ_μ_*(*x,t*). In our analysis below, we will assume that the ecosystem has approached a steadystate, in which the species densities *ρ_μ_*(*x, t*) and environmental variables 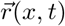 can be described by time invariant profiles *ρ_μ_*(*x*) and 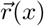. This is a crucial simplification, as it means that the growth rate 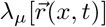 can be replaced by a simple spatially varying growth function 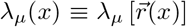, which is conditionally independent of the surrounding community. The steady-state growth profiles are uniquely determined by the steady-state density profiles,

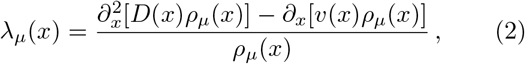

such that the realized growth rates can in principle be determined from static snapshots of a community’s spatial composition.

**FIG. 1.**
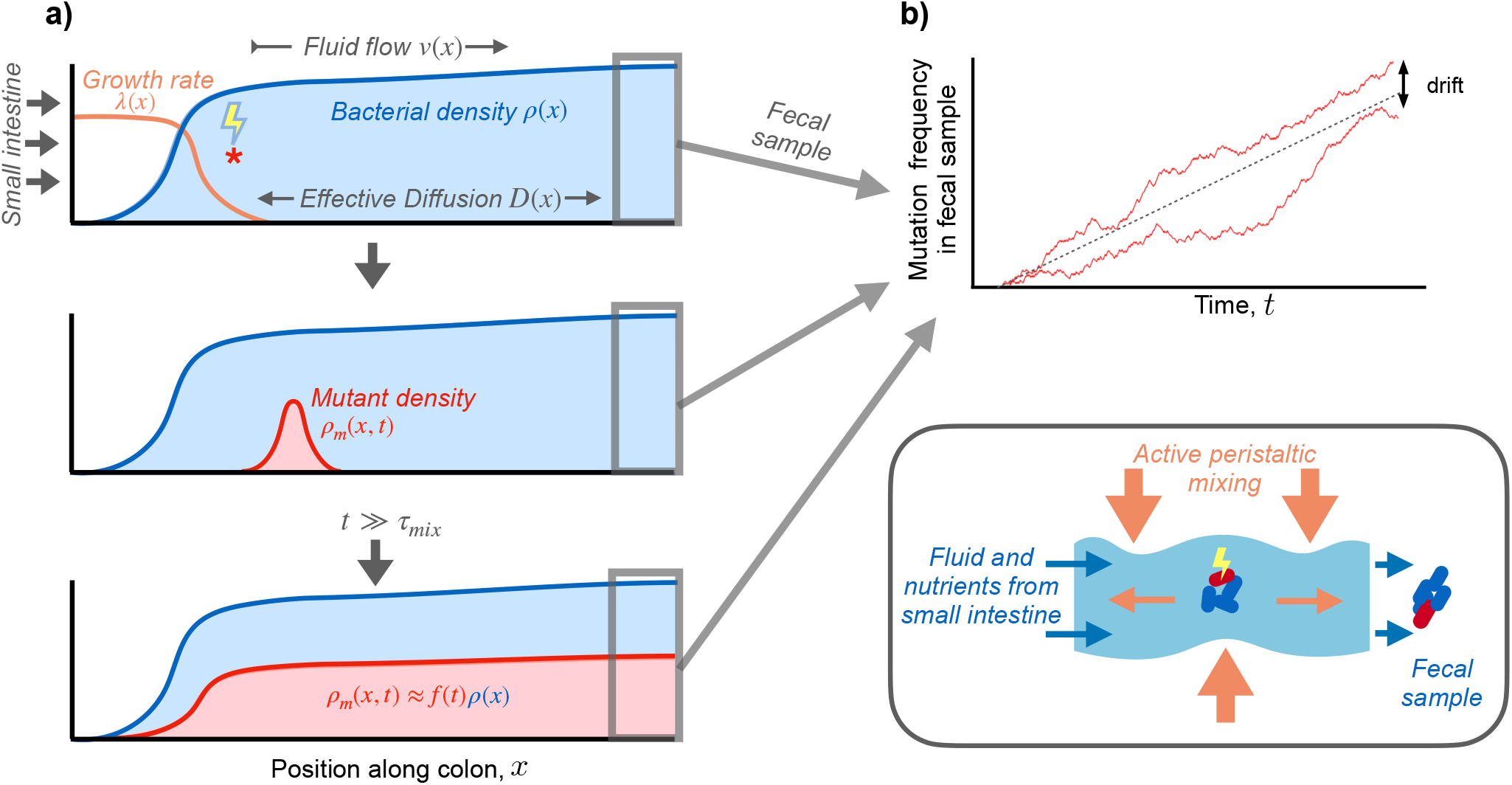
Dynamics of a genetic variant in a spatially extended population in the intestinal lumen. (a) Schematic of spatial growth model. Nutrients enter the colon from the small intestine (left) and fecal samples are generated at the distal end (right). Intestinal wall contractions mix contents of lumen (inset), producing effective diffusion in the axial direction (26, 28). Continuous fluid flow through the colon generates longitudinal gradients in steady-state bacterial growth rates [λ(*x*), orange line] and population densities [*ρ*(*x*), shaded blue region]. Mutation events (red star) found new lineages that begin to grow and spread out in space [*ρ_m_*(*x,t*), shaded red region]. After Tmix generations, the spatial distribution of the mutant lineage approaches that of the wildtype population, but with a smaller overall size. (b) Mutation frequencies observed in fecal samples over time. Two hypothetical trajectories are shown (red lines), illustrating the stochastic fluctuations that arise due to genetic drift.

### Dynamics of mutant lineages

We wish to understand the dynamics of genetic variants within species in the steady-state communities described above. Since these variants may initially be present in just a small number of cells, the deterministic dynamics in Eq. (1) must be augmented to account for stochastic fluctuations in cell numbers in both space and time. As long as the frequency of the mutant lineage is sufficiently rare, we expect it to have a negligible influence on the steady state values of *r_i_*(*x*) and *ρ_μ_* (*x*), such that the growth function of the mutant can be described its own steady state growth profile *λ_m_* (*x*). In Supplementary Information 1, we use a microscopic birth-and-death model to show that the density profile of the mutant lineage can be described by the coarse-grained stochastic differential equation

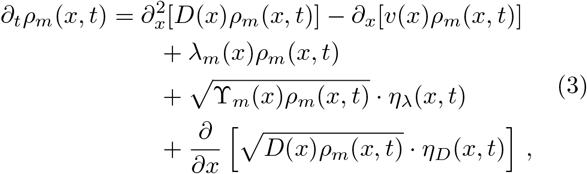

where *η_λ_*(*x,t*) and *η_D_*(*x,t*) are independent Brownian noise terms with 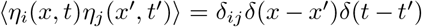 (43), and 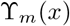 is the variance in offspring number of the local birth and death process. In the absence of cell death, 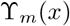 will be proportional to the local growth rate *λ_m_* (*x*); we expect this to be a reasonable approximation in the human gut, where live cells comprise a large fraction of the total excreted biomass (5, 44, 45).

Equation (3) is a stochastic variant of the familiar Fisher-KPP equation (46, 47), which is frequently used to model the spread of species or genetic variants through space (46, 47, 48, 49). An important difference in our setting is that the expansion velocity *v*(*x*) is now set by the fluid flow in the environment, such that the mutant and wildtype populations are continuously “treadmilling” in place. We will show below that this typically leads to “pushed” rather than “pulled” expansions (50, 51), which will have important implications for the stochastic dynamics of genetic drift in this system (52, 53).

Equation (3) provides a quantitative model for predicting the spread of a genetic variant within a resident gut community under the combined action of fluid flow, mixing, and spatially varying growth rates. While the spatiotemporal dynamics of Eq. (3) are often quite complicated, further simplifications arise in scenarios where the emergent evolutionary forces (e.g. natural selection and genetic drift) are much slower than the “ecological” processes of growth, flow, and mixing. These ecological forces will lead to a characteristic *mixing timescale τ*_mix_ (discussed in more detail below), which represents the time required for the descendants of a mutant lineage to disperse across space (Fig. 1A). For times much larger than *τ*_mix_, the shape of the mutant density profile will approach the steady-state population profile,

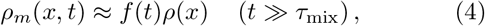

but with a prefactor *f* (*t*) that depends on the growth rate λ_*m*_(*x*), the stochastic noise terms in Eq. (3), and the initial density profile *ρ_m_*(*x*, 0). This prefactor *f* (*t*) also coincides with the relative frequency of the mutant lineage in later fecal samples (Fig. 1B). Thus, the dynamics of *f* (*t*) provide a crucial connection between the spatial growth dynamics in Eq. (3) and the frequencies of mutant lineages that are measured in metagenomic sequencing data.

Much of the relevant behavior can be understood by considering a perfectly neutral lineage [λ_*m*_(*x*) = λ(*x*)] founded by a single individual a position *x* (Fig. 1A). In Supplementary Information 2, we show that the distribution of *f* (*t*) at intermediate times (*t* ≫ *τ*_mix_) can be captured by a well-mixed Wright-Fisher model

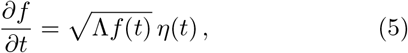

with an effective initial frequency

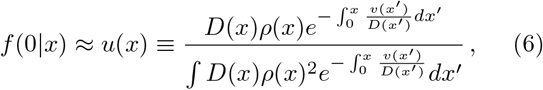

and an effective rate of genetic drift

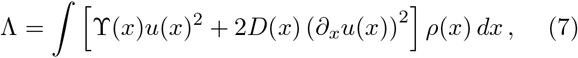

which is independent of the mutation’s initial position. Equation (5) implies that this new mutation will require a time of order *τ*_drift_ ~ *f*/Λ to drift to frequency ~*f* in future fecal samples (54, 55). These dynamics will be consistent with the separation-of-timescales assumption as long as Tdrift *τ*_mix_, which requires that *f* (*t*) ≫ Λ*τ*_mix_. Since Λ is inversely proportional to the bacterial density, and *τ*_mix_ is independent of density (see below), we expect these results to apply for a broad range of frequencies when population sizes are sufficiently large.

At intermediate times (*τ*_mix_ ≪ *t* ≪ 1/Λ), the descendants of the mutant lineage will be approximately evenly distributed across the population (Fig. 1A), so the conditional fixation probability of the entire lineage is simply *f* (*t*). Averaging over the random values of *f* (*t*) using Eq. (5), we find that

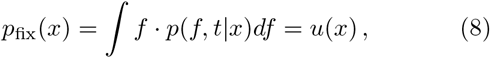

so that Eq. (6) has a simple interpretation as the fixation probability of a neutral mutation at position *x* (37). Similarly, the product *g*(*x*) = *u*(*x*)*ρ*(*x*) represents the spatial distribution of future common ancestors of the population (37) — regardless of whether they have a mutation or not — which provides a quantitative measure of the spatial locations that have an outsized influence on the genetic composition of future fecal samples. By modelling the long-term accumulation of neutral mutations within these populations, we obtain a simple prediction for the effective generation time,

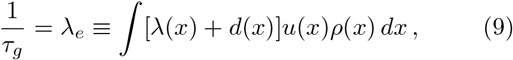

where *d*(*x*) is the local death rate (Supplementary Information 2.3). This allows us to re-express the “wall-clock” rate of genetic drift in Eq. (7) as a more traditional effective population size:

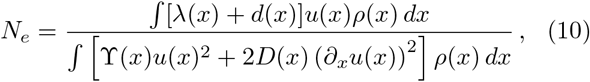

which is proportional to the total bacterial density as expected. When cell death is negligible 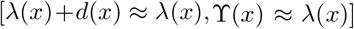, these effective parameters are completely determined by the steady state density profile *ρ*(*x*).

### Analytical solutions for a minimal model of bacterial growth

We can understand the implications of these results by examining a minimal model of bacterial growth in the gut, which yields analytical solutions for the growth and density profiles in Eqs. (9) and (10). We assume that flow and diffusion are uniform in space, such that there is a characteristic length scale ~D/v that sets the transition between diffusive and advective motion. We further assume that growth depends on a single limiting nutrient with a step-like dose response curve (Fig. 2A). These assumptions produce a spatial growth profile with a sharp transition at a critical position *x* = *ℓ* (Fig. 2B), whose location depends on the relative magnitudes of λ, *v*, and *D*. In Supplementary Information 4, we obtain analytical expressions for the resulting population density profiles, which are illustrated in Fig. 2C,D.

**FIG. 2.**
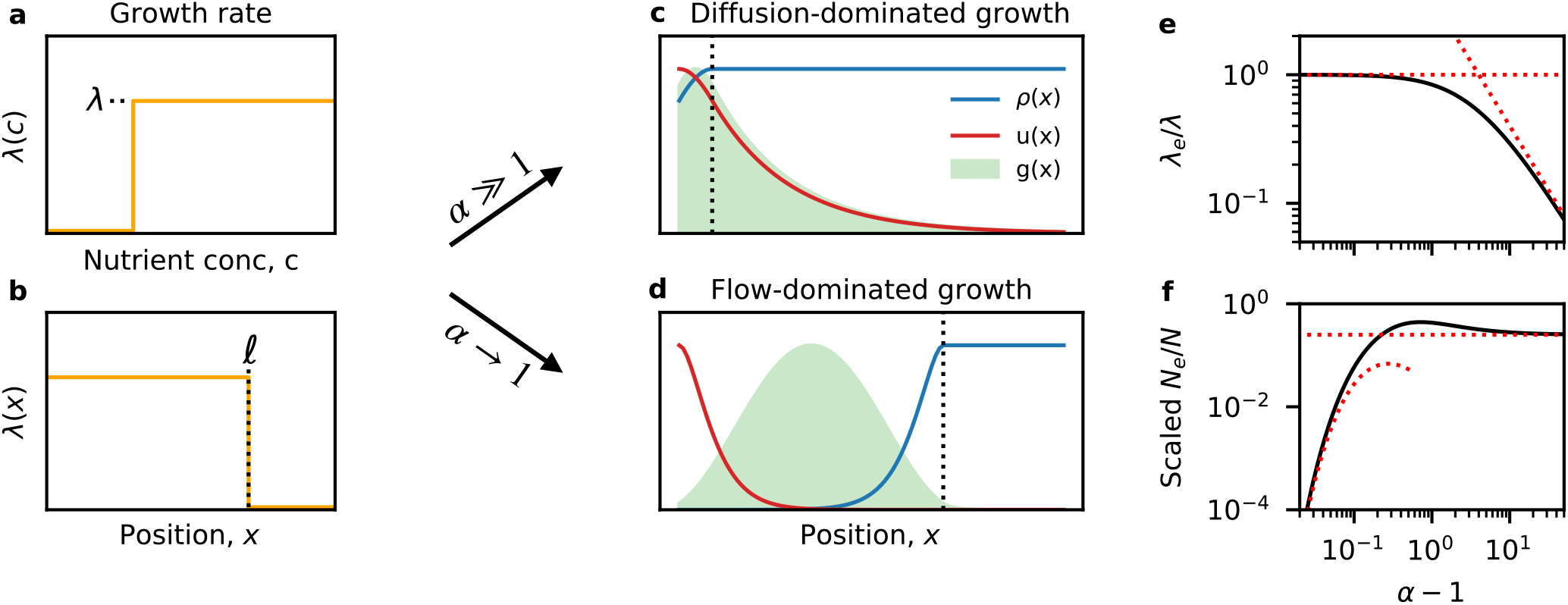
Analytical solutions for a minimal model of bacterial growth. (a,b) Schematic of bacterial growth rates. A step-like growth response curve (a) leads to a sharp transition between high growth (λ(*x*) ≈ λ) and no-growth (λ(*x*) ≈ 0) at a critical position *ℓ* (b). (c, d) Spatial profiles of population density, *ρ*(*x*) (blue line), neutral fixation probability, *u*(*x*) (red line), and distribution of future common ancestors, *g*(*x*) = *u*(*x*)*ρ*(*x*) (green shaded region), for examples of (c) diffusion-dominated growth (4*D*λ/*v*^2^ ≈ 10) and (d) flow-dominated growth (4*D*λ/*v*^2^ ≈ 1.04). In both cases, the dashed black lines indicate the boundary of the growth region in panel b. (e) Scaled effective growth rate, λ_*e*_/λ, and (f) scaled effective population size, (*N_e_/N*} · (*vL*/2*D*), as a function of the washout parameter *α* ≡ 4*D*λ/*v*^2^. Black lines denote the analytical predictions in Supplementary Information 4.2, while the red lines illustrate the asymptotic behavior in the *α* 1 and *α* – 1 ≪ 1 limits.

We observe two characteristic regimes that depend on the value of the dimensionless parameter *α* ≡ 4*D*λ/*v*^2^, which compares the characteristic timescales of growth (λ^-1^) and local dilution (~*D/v*^2^). When *α* < 1, growth is not sufficient to overcome the constant loss of individuals due to fluid flow, and the population cannot permanently establish itself in the gut — a scenario we refer to as “washout” (26). At the opposite extreme (*α* ≫ 1), growth is sufficiently rapid that the nutrients are depleted long before flow is relevant (*ℓ* ≪ *D/v*), and diffusive mixing is still the dominant form of motion. This diffusion-dominated growth produces a nearly uniform population density profile, and a distribution of future ancestors [*g*(*x*) = *u*(*x*)*ρ*(*x*)] that is primarily composed of slowly growing cells (Fig. 2C). This leads to a reduced effective growth rate, λ_e_~ *v*^2^/*D* ≪ λ (Fig. 2E), and an effective population size, *N_e_*~*ρ*(*L*)*D/v*, that is independent of the length of the colon (Fig. 2F).

Between these two extremes, we also observe an intermediate regime close to the washout threshold (*α* → 1), where flow dominates over diffusion throughout most of the growth region (*D/v* ≪ *ℓ*). This produces a strong spatial gradient in the density profile, and a distribution of future ancestors [*g*(*x*) = *u*(*x*)*ρ*(*x*)] that is primarily concentrated within the growth region (Fig. 2D). The effective growth rate therefore approaches its maximum value λ_*e*_ ≈ λ (Fig. 2E), while the effective population size is dramatically reduced (Fig. 2F). By examining the contributions to Eq. (10), we find that these lower effective population sizes are driven by fluctuations in a narrow region near the entrance of the colon (*x* ≾ *D/v*), rather than the comparatively larger sub-population of dividing cells (*x ~ ℓ*). This shows how long-term lineage dynamics can have a major impact on the emergent evolutionary parameters in our model.

While the enhanced fluctuations are most pronounced near the washout threshold (*α* ≈ 1), we note that washout itself does not play a major role in this behavior. In Supplementary Information 4.3, we show that the results in Fig. 2 are robust to the addition of a washed out region at the proximal end of an otherwise established population. This suggests that it is the accumulated growth over many diffusion lengths — rather than washout per se — that is the primary cause of the reduced effective population size when *α* → 1. It also suggests that our qualitative results are do not depend on the specific nature of the boundary between the large and small intestine: the small number of cells that diffuse into this low-growth region do not dramatically change the dynamics that occur downstream.

### Dynamics of selected mutations

We can use a similar separation-of-timescales approach to analyze the fates of selected mutations as well (Supplementary Information 3). Provided that the growth rate differences are sufficiently small (see below), we find that the emergent dynamics on timescales *t* ≫ *τ*_mix_ can again be mapped on to well-mixed Wright-Fisher model

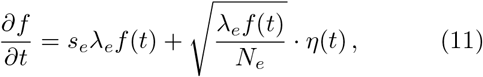

with an effective selection coefficient

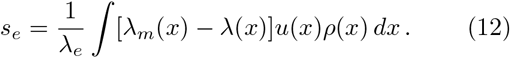

This equation encapsulates the relevant growth rate differences across the ancestral region *g*(*x*) ∝ *u*(*x*)*ρ*(*x*). For a global increase in growth rate [λ_*m*_(*x*) = (1 + *s*)λ(*x*)], the effective selection coefficient *s_e_* is proportional to s as expected. However, Eq. (12) also allows us to account for tradeoffs in growth rate that emerge at different spatial locations. The relative weights of fitness effects in these different microenvironments are determined by the product λ(*x*)*g*(*x*), which biases the effective selection coefficients toward regions with higher local growth rates.

The mapping in Eq. (11) allows us to obtain a precise picture of the dynamics a beneficial mutation as observed in sequenced fecal samples. When *τ*_mix_ ≪ *t* ≪ 1/*s_e_*λ_*e*_, the dynamics of the selected lineage will be identical to the neutral mutations described above. With probability ~*N_e_s_e_u*(*x*), the mutant lineage will eventually establish and grow deterministically as ~*e*^*s*_*e*_λ_*e*_*t*^, and will fix within the population on a characteristic timescale 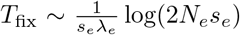 (54, 55). These dynamics will be consistent with the separation-of-timescales assumption as long as *s_e_*λ_*e*_*τ*_mix_ ≪ 1.

For stronger selection strengths (*s_e_*λ_*e*_*τ*_mix_ 1), the separation-of-timescales assumption will eventually break down, and growth rate differences will start to play an important role during the mixing process itself. In this case, we show in Supplementary Information 3.1 that an analogous fixation probability can still be obtained numerically by solving the ordinary differential equation

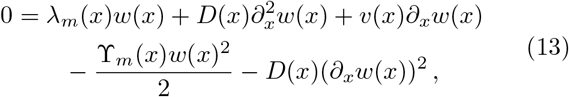

which gives the fixation probability of a beneficial mutation that arises at a given position x. Equation (13) is similar to existing branching process models of fixation in spatially or genetically subdivided populations (56, 57, 58), but with an additional non-linearity that accounts for the conservative noise of spatial diffusion. The total fixation probability can be obtained from Eq. (13) by averaging *w*(*x*) over the local production rate of new mutations, *μ*(*x*)*ρ*(*x*), yielding

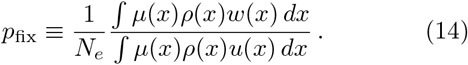

When *s_e_*λ_*e*_*τ*_mix_ ≪ 1, we recover the linear relation *p*_fix_ = 2*s_e_* predicted by the separation-of-timescales approximation. Deviations from this behavior at larger values of *s_e_* can therefore provide insight into the mixing timescale *τ*_mix_, which we have so far treated as an implicit parameter.

In Fig. 3, we compute these fixation probabilities numerically for the minimal model in Fig. 2, assuming a global increase in growth rate [λ_*m*_(*x*) = (1 + *s*)λ(*x*)] and a replication-dominated mutation rate [*μ*(*x*) ≈ *μ*λ(*x*)]. We find good agreement with the separation-of-timescales prediction at sufficiently low values of *s*. This shows that *τ*_mix_ is primarily determined by the ancestral region [*g*(*x*) = *u*(*x*)*ρ*(*x*)] rather than the total length of the colon (*L* → ∞). For diffusion-dominated growth (4*D*λ/*v*^2^ > 1), this linear approximation remains accurate for growth rate advantages as large as *s*~100%, demonstrating the broad utility of the separation-of-timescales approach in this regime. In contrast, we find that deviations can emerge at much lower values of *s* in the flow-dominated growth regime (4*D*λ/*v*^2^ – 1 ≪ 1), such that the fixation probabilities are often much larger than expected under the separation-of-timescales approximation (Fig. 3C). The spatial fixation profiles in Fig. 3B show that this increase is primarily driven by enhanced fixation probabilities in lower regions of the colon (Fig. 3C), which acts to expand the range of production of successful beneficial mutations. These deviations can be rationalized by the longer times required for lineages to migrate across the ancestral region ~*v/ℓ*, such that even small growth rate advantages can produce large numbers of additional offspring before complete mixing is achieved. However, despite these enhanced fixation probabilities, we find that the long-term growth rates of established mutations continue to match the separation-of-timescales prediction above. By solving a related equation for the average density profile (Supplementary Information 4.5), we find that the exponential growth rates eventually approach *S* ~ *s_e_*λ_*e*_ even when *s_e_*λ_*e*_*τ*_mix_ ≫ 1. This shows that finite mixing can have a differential impact depending on the timescales of the corresponding evolutionary processes (e.g. establishment vs fixation).

**FIG. 3.**
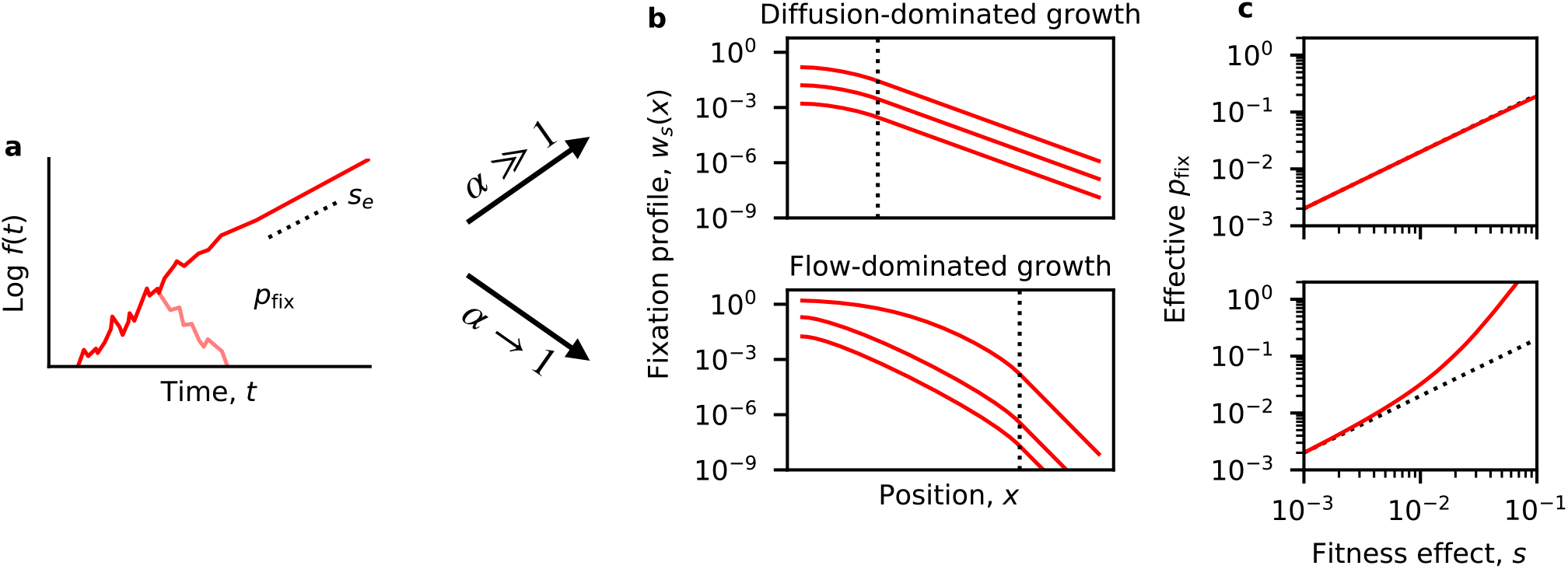
Dynamics of selected mutations. (a) Schematic trajectory of a beneficial mutation, which either establishes and grows exponentially at rate *s_e_* (dashed line) or else goes extinct (light red line). (b) The spatially-resolved fixation probability *w_s_*(*x*) of a beneficial mutation that originates at position *x*. Red lines denote the numerical solutions of Eq. (13) for the minimal model in Fig. 2 with a constant growth rate advantage λ_*m*_(*x*) = (1 + *s*)λ(*x*) (Supplementary Information 4.4). The dashed lines indicate the transition from λ(*x*) ≈ λ to λ(*x*) ≈ 0. The top panel illustrates a population in the diffusion-dominated growth regime (4*D*λ/*v*^2^ ≈ 1.9) while the bottom panel illustrates a population in the flow-dominated growth regime (4*D*λ/*v*^2^ ≈ 1.04). In both cases, fixation profiles were calculated for *s* = 10^-3^, 10^-2^, and 10^-1^. (c) The effective fixation probability obtained by averaging *w_s_* (*x*) the range of initial positions. The top and bottom panels denote the same populations as in (b). Dashed lines indicate the separation-of-timescales prediction, *p*_fix_ = 2*s_e_*. Deviations from this prediction emerge in the flow-dominated growth regime, due to the enhanced fixation probability of mutations that originate at larger *x* values (illustrated in panel b).

## APPLICATION TO THE HUMAN COLON

Our analytical results have shown that the effective evolutionary parameters can strongly depend on the relative magnitudes of growth, flow, and mixing across the large intestine. How do these results extend to bacteria growing in the human colon? While direct measurements of the growth rates and density profiles are difficult to obtain in this system, rough estimates for many of the key parameters were recently obtained by Cremer *et al* (28) using a generalization of the model in Eq. (1). By explicitly modeling the consumption and production metabolites along the length of the human colon (*L*~2*m*), Cremer *et al* (28) obtained predictions for the steady-state growth rates and density profiles for a pair of representative species from the *Bacteroides* and *Firmicutes* phyla (Fig. 4A).

**FIG. 4.**
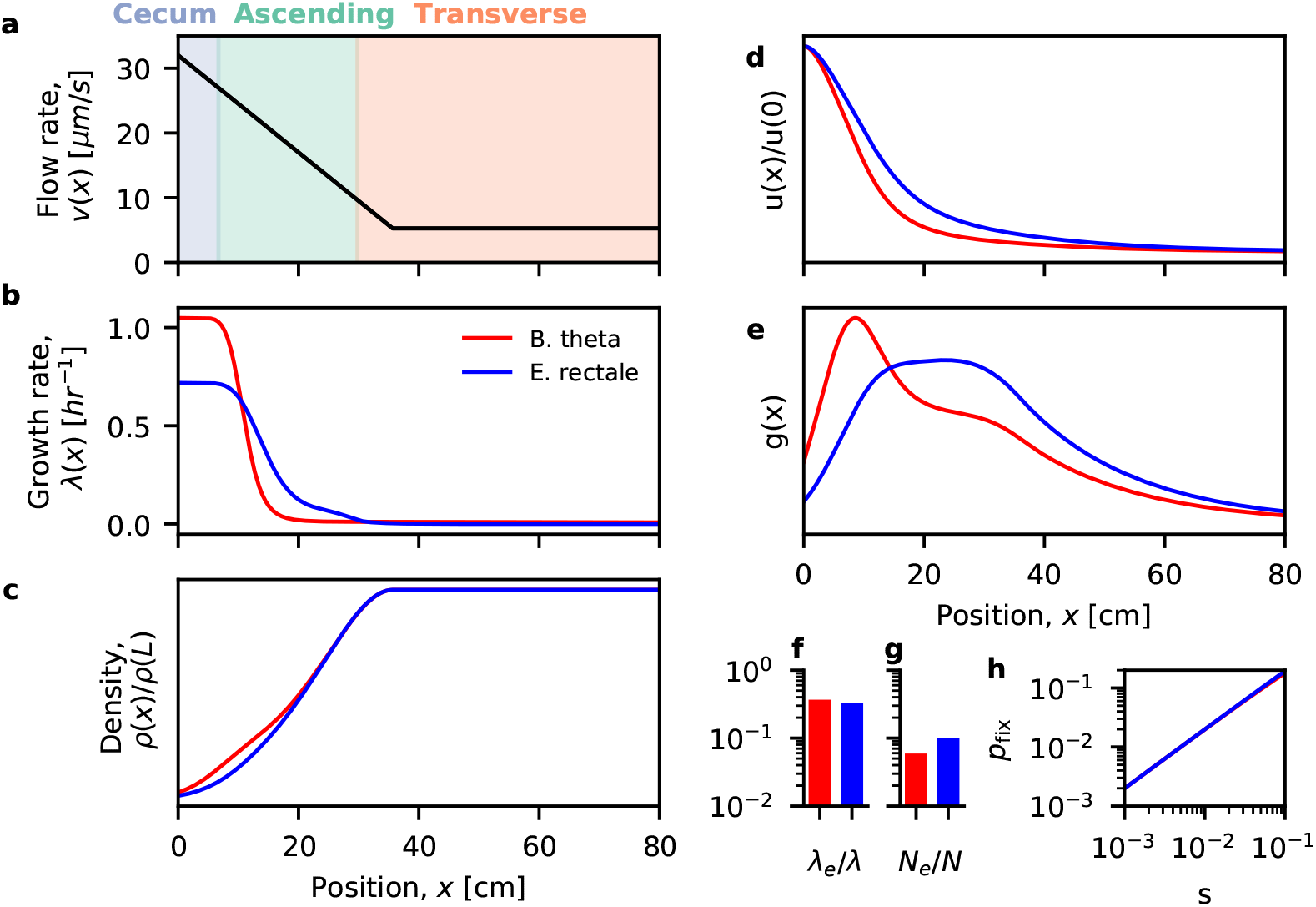
Applications to bacteria in the human colon. (a) Estimates of the flow velocity along the first 80 cm of the human colon (28). Colors indicate the positions of different colonic segments. (b,c) Steady-state growth rate and density profiles estimated for two representative species using the results of Ref. (28). Growth rate profiles were extracted from the figures in Ref. (28), and the corresponding density profiles were calculated from Eq. (2) using the methods described in Supplementary Information 5. (d,e) Predictions for the neutral fixation profile u(x) (d) and distribution of common ancestors g(x) (e) obtained from the estimated parameters in (a-c) (Supplementary Information 5). In both cases, the predicted ancestral distributions contain a large contribution from regions with low growth rates. (f-g) Predictions for the effective growth rate, effective population size, and the net fixation probability of beneficial mutations (Supplementary Information 5).

An important feature of this model is a spatially varying flow rate, due to the absorption of water by the colonic epithelium. Cremer *et al* (28) estimated that this effect causes the flow rate to decline from *v* ≈ 30 *μm/s* to *v* ≈ 5 *μm/s* over the length of the proximal colon (≈30cm). Given an estimated mixing rate of *D*≈10^6^*μm*^2^/*s*, the effective diffusion lengthscales range from *D/v* ≈ 3 — 20*cm*, and the corresponding timescales range from *D/v*^2^≈16 minutes to 11 hours. These estimates suggest that the *in vitro* growth rates of the representative strains (λ = 0.6 – 1 hr^-1^) will be close to washout at the highest flow rates, but far from washout elsewhere, making it difficult to directly apply the results from our minimal model in Figs 2 and 3.

To overcome this challenge, we developed a numerical approach for calculating the effective evolutionary parameters in Eqs. (9), (10), and (14) using the steadystate density profiles estimated by Cremer *et al* (28) (Supplementary Information 5). The results are shown in Fig. 4. In both cases, we find that the flow rate decreases sufficiently rapidly across the proximal colon that the corresponding ancestral regions [*g*(*x*) = *u*(*x*)*ρ*(*x*)] include a substantial contribution from slowly growing cells. For example, the *B. theta* curves in Fig. 4E predict that about a third of the future common ancestors will come from regions where growth rates are <0.5/day, while another third will come from nutrient-rich regions where growth rates are >0.5/hr. This suggests that these resident populations will lie on the boundary between diffusion-dominated and flow-dominated growth in Figs 2 and 3 above. This has several implications for the emergent evolutionary dynamics:

Figure 4F shows that the effective growth rates are about 3 times smaller than the maximum growth rates attained at the entrance of the colon. This corresponds to about 9 generations per day for the *B. theta* example and about 5 generations per day for *E. rectale.* These predictions are 5-10 times higher than the traditional ~1/day estimates based on the daily biomass excreted in feces (59, 60), which reflects the higher-than-average growth rates experienced by evolutionarily relevant lineages. Interestingly, these higher growth rates are more consistent with direct measurements of *B. theta* generation times in germ-free mice (61), suggesting that some of the emergent evolutionary parameters may be similar in these environments, despite the vast differences in host physiology.

In addition to these shortened generation times, Fig. 4G shows that corresponding effective population sizes are about 10-20 times smaller than the total number of bacterial cells in the colon. This leads to elevated rates of genetic drift, but not at the levels necessary to drive large changes in the frequencies of genetic variants over a single host lifetime. Given the large number of bacteria in the human gut [*N* ~ 10^13^ (62)], these estimates predict that a species with 1% relative abundance would have an absolute rate of genetic drift of about Λ≡λ_*e*_/*N_e_*~10^-9^ per day. At this rate, genetic drift would require more than a hundred years to generate even a 1% shift in the frequency of a common genetic variant. This stands in contrast to the rapid shifts in frequency that are observed for some mutations in sequenced human fecal samples (Fig. 5). This quantitative discrepancy suggests that other factors, like natural selection or temporal bottlenecks, are required to explain these observations.

**FIG. 5.**
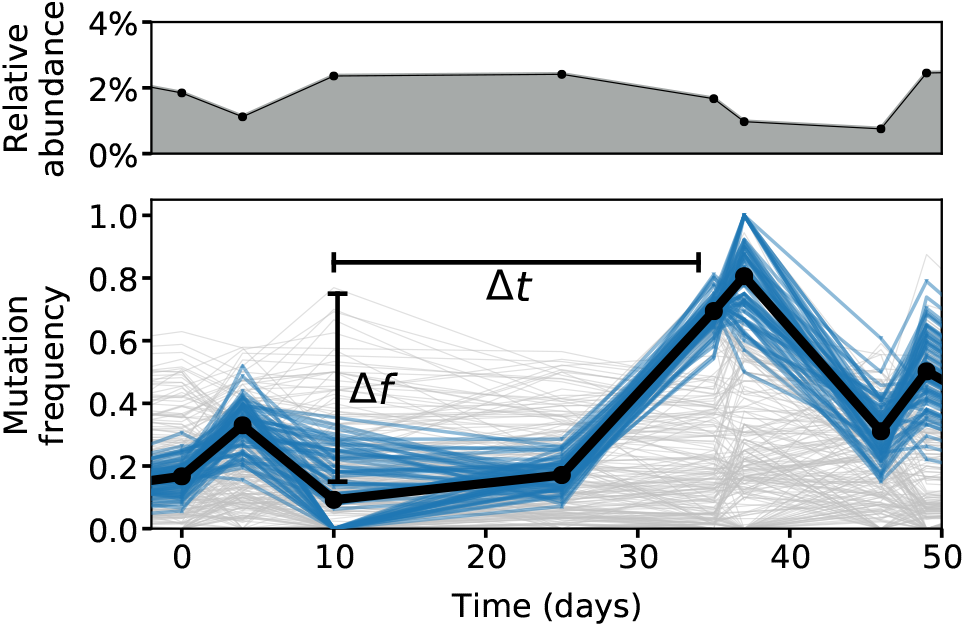
Mutation frequencies within a *Bacteroides massiliensis* population from a single human subject. Relative abundance of *B. massiliensis* (top) and the frequencies of single nucleotide variants in this species (bottom) inferred from longitudinally sequenced fecal samples from a previous study (15) (Methods). Blue trajectories highlight a cluster of mutations that experienced a large shift in frequency during the study (Δ*f* ≈ 50%), despite relatively small changes in the overall relative abundance of **B.** *massiliensis* (~2-fold; top). The timescale of this genetic shift (Δ*t* ~ 25 days) is many orders of magnitude faster than our predicted timescale of genetic drift (Δ*t* ~ 1/Λ; Fig. 4 and Eq. 16), even after accounting for the observed fluctuations in overall relative abundance.

Finally, Fig. 4H shows that the mixing timescales in the human colon are sufficiently rapid that the establishment probabilities of beneficial mutations are well-approximated by the linear prediction [*p*_fix_~2*s_e_* across a wide-range of physiologically relevant fitness benefits. These results show that our separation-of-timescales approach can apply even for human-relevant conditions, yielding quantitative estimates for a number of key evolutionary parameters.

### Extensions to wall growth and other spatial refugia

So far, we have assumed that the emergent evolutionary forces are dominated by the dynamics within the luminal fluid. However, our basic mathematical framework can also be extended to incorporate certain forms of wall growth (or other spatial refugia), which are known to play an important role in a variety of microbial ecosystems (23, 25, 42, 63, 64). While a complete analysis of this case is beyond the scope of the present paper, many of the key features can already be appreciated within a simple model, in which a well-mixed luminal compartment is connected to a large number of protected sub-populations on the walls of the large intestine (Fig. S1, Supplementary Information 6). By mitigating the effects of fluid flow, these wall-bound populations have an advantage in contributing genetic material to the future population. In extreme cases, these wall-bound populations can help recolonize the lumen after strong perturbations like diarrhea. However, their influence on the steady-state dynamics of genetic turnover in the gut are less well understood.

Previous work has argued that wall growth comprises a small fraction of the daily biomass production in the human gut, since the effective linear density in the wall (*ρ_w_*) is much smaller than the corresponding density in the lumen (*ρ_ℓ_*) (26, 28). By generalizing our separation-of-timescales approach to this regime, we find that the longterm rates of genetic turnover critically depend on the *engraftment rate ε*, which represents the per-generation probability for a luminal cell to successfully migrate into one of the subpopulations on the wall (Supplementary Information 6). When wall growth comprises a small fraction of the total biomass production (*ρ_w_*λ_*w*_ ≪ *ρ_ℓ_*λ_*ℓ*_), we find even tiny rates of engraftment,

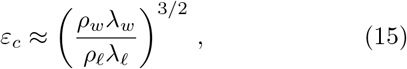

are sufficient to ensure that genetic drift is dominated by the luminal compartment (Λ ≈ Λ_*ℓ*_). This suggests that there is a broad parameter regime where our previous results still apply. When *ε* ≪ *ε_c_*, fluctuations within the wall population start to become more important, and the long-term rate of genetic drift is amplified by a factor of

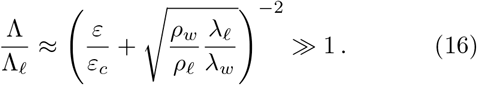

At even smaller engraftment rates, the separation-of-timescales approximation eventually breaks down, and the long-term dynamics start to be dominated by the sequential spread of strains across many isolated refugia (Supplementary Information 6). These slow spreading dynamics can lead to apparent rates of genetic drift that are even weaker than the lumen (Λ ≪ Λ_*ℓ*_), despite the fact that the relevant population of cells is significantly smaller. This highlights how the dynamics of many distinct refugia can be qualitatively different than a single refuge on its own.

This non-monotonic behavior illustrates the importance of the engraftment rate *ε*, which is currently poorly constrained by experimental data. Previous estimates in the human gut suggest that the total wall growth is bounded by *ρ_w_*λ_*w*_/*ρ_ℓ_*λ_*ℓ*_ ≾ 10^-3^ (26). Using these estimates as a guide, our model predicts that wall growth should have a negligible impact on genetic drift as long as *ε* ≿ 10^-4^, and that engraftment rates of order *ε*~10^-6^. are required to enhance the rate of drift by a factor of ~10^3^. These values are still too small to explain the rapid shifts in frequency observed in sequenced fecal samples (Fig. 5).

Our analysis also highlights the importance of the underlying timescales of these dynamics: while smaller engraftment rates can produce higher rates of genetic drift, these fluctuations still require at least ~*ρ_ℓ_*λ_*ℓ*_/*ρ_w_*λ_*w*_ generations to propagate back into the lumen before they are eventually sampled in feces (Supplementary Information 6). On shorter time scales, we expect that wall-dominated genetic drift will generate autocorrelated fluctuations in sequenced fecal samples, similar to the background selection scenario analyzed in Ref. (40). Understanding the dynamics of these lagged fluctuations – and the signatures they leave in genetic data like Fig. 5 – could provide a promising approach for measuring the impact of wall growth from bulk metagenomic sequencing data.

## DISCUSSION

The spatial organization of microbial communities can strongly influence their ecological and evolutionary dynamics. Here we sought to quantify these effects in the human gut microbiota, whose long-term composition is shaped by lineages that manage to evade eventual dilution in feces. We have developed a mathematical framework for predicting how stochastic evolutionary forces emerge from simple models of bacterial growth in the intestinal lumen, and how they give rise to the genetic variation that can be observed in sequenced fecal samples.

While our simplified models neglect many important features of microbial and host physiology, they make several new predictions about the role of spatial structure for the within-host evolution of the gut microbiota. Chief among these is the finding that the emergent evolutionary dynamics can often be captured by effective models that lack explicit spatial structure, even in cases where longitudinal gradients have a large effect on local species composition. Similar results are implicit in earlier studies that focused on neutral fixation and heterozygosity (37, 38, 39, 65). Here we showed that this equivalence extends to wider range of evolutionary phenomena, including the establishment and spread of beneficial mutations. This suggests that well-mixed models from population genetics might have broader utility for modeling the dynamics of genetic variants in sequenced fecal samples, despite their explicit spatial structure. In these cases, our results provide a basis for connecting the inferred parameters (e.g. effective selection coefficients and population sizes) with specific weighted averages over the underlying growth and density profiles.

Consistent with previous intuition, we found that unidirectional fluid flow can have a large impact on the emergent evolutionary parameters in the gut. Populations that are close to the washout threshold experience dramatically elevated rates of genetic drift — much larger than would be predicted by the absolute number of dividing cells (Supplementary Information 4.2). This shows how long-term lineage dynamics can play an essential role in extrapolating evolutionary parameters from absolute growth and density measurements.

By applying our results to recent estimates from the human gut, we found that continuous fluid flow (and simple forms of wall growth) are unlikely to lower effective population sizes to the point where we would expect large fluctuations in genetic composition within a host lifetime. This suggests that additional factors, such as natural selection or temporal bottlenecks, are necessary to explain the rapidly changing mutation frequencies that have been observed in sequenced human fecal samples (7, 13, 14, 15, 16, 17, 18, 19, 20). We also found that the spatial gradients generated by fluid flow can cause the effective generation time to be about 5-10 times shorter than the traditional ~1/day estimates obtained from the average growth rate across the colon (59, 60). This suggests that the human gut microbiota may experience substantially more evolution than previously expected over the course of a given year. It also suggests that relatively small fitness advantages (*s*~1 —10%) may be sufficient to drive large changes in the frequencies of genetic variants over daily and weekly timescales. We note that our precise estimates should be treated with a degree of caution, since they are currently derived from relatively rough estimates the underlying growth rates and density profiles in the gut (28). Our framework provides a flexible approach for refining these estimates using future measurements of spatially resolved growth rates (66) or microbial abundances (32) in human subjects. Our results also highlight the spatial locations where additional measurements would be most informative for constraining our estimates of the emergent evolutionary parameters.

Future experiments could test these predictions quantitatively using genetic barcoding techniques (67, 68, 69), both in animal models (12), and potentially even in humans (70). For example, by introducing libraries of barcoded strains and tracking their relative fluctuations in subsequent fecal samples, it should be possible to directly measure the effective rate of genetic drift in Eq. (5) to test our theoretical predictions in Fig. 4. Even stronger tests are possible if the strain labeling events are coupled to local environmental conditions (e.g. pH, mucin, etc.) (71, 72, 73). By tracking the frequencies of these labeled lineages as they spread out across the colon — and potentially into the intestinal walls (32) — one could directly measure the weighting function *u*(*x*) and its associated mixing timescales. Our results provide a framework for interpreting these spatiotemporal measurements, and for using them to extract key unknown parameters like the engraftment rate *ε* (Supplementary Information 6).

The limitations of our simple growth model create many additional directions for future work. For example, our results suggest that it will be critical to understand how temporal variation in growth conditions – either from regular circadian rhythms (74, 75) or larger perturbations like antibiotics (15) – might alter the emergent evolutionary forces we have studied here. It would also be interesting to examine how more complex forms of wall growth or other spatial refugia (61, 76, 77) might contribute to long-term genetic turnover within the gut. While the details of these models will necessarily differ from the ones studied here, our results show how similar separation-of-timescales approaches may continue to be useful for analyzing these additional scenarios as well. By “integrating out” variation over shorter length- and time-scales, these coarse-graining approaches can provide a promising route for modeling evolutionary dynamics in complex natural settings (78).

## DATA AND CODE AVAILABILITY

Source code for the data analysis and figure generation is available at Github (https://github.com/bgoodlab/spatial_gut_evolution).

## ACKNOWLEDGMENTS

This work was supported in part by a National Science Foundation Graduate Research Fellowship, the Alfred P. Sloan Foundation, and a Terman Fellowship from Stanford University.

## SUPPLEMENTAL INFORMATION (SI)

### 1. MICROSCOPIC DERIVATION OF THE REACTION-DIFFUSION MODEL

In this section, we introduce a microscopic birth-and-death model that reproduces the continuum limit described by Eq. (3). We begin by deriving a mean-field model for the total population profile in Eq. (2). We assume that space and time are divided into discrete intervals of size Δ*x* and Δ*t*, respectively. In a given interval Δ*t*, we will let *m_v_* (*x*) denote the probability that an individual is carried from position *x* to position *x* + Δ*x* by advection (flow). Likewise, we will let 2*m_D_* (*x*) denote the probability that an individual moves to either *x* + Δ*x* or *x* – Δ*x* through diffusion (mixing). Finally, we will let *m_b_*(*x*|{*ρ*(*x, t*)}) and *m_d_*(*x*|{*ρ*(*x, t*)}) denote the probability that an individual gives birth or dies, respectively; these probabilities will generally depend on both the position *x* as well as the current density profile, *ρ*(*x, t*). In the limit of large densities, these assumptions yield a mean-field update rule:

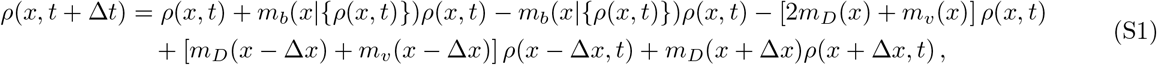

whose continuum limit is given by

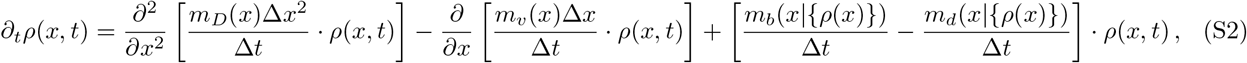

This allows us to identify the macroscopic parameters

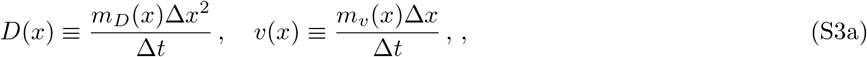

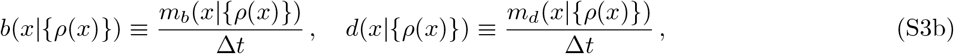

as well as the net growth rate,

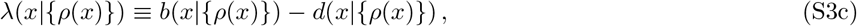

so that

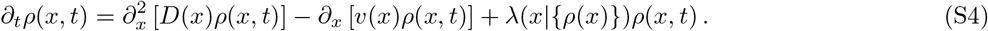

At steady state, this reduces to

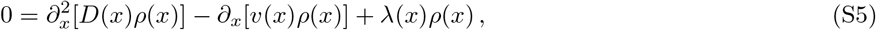

which can be rearranged to obtain Eq. (2) in the main text.

The relevant boundary conditions can be derived in several different ways. In the simplest version, we assume that there is no input or output flux of bacteria from advection or diffusion through the left boundary (*x* = 0). This implies that

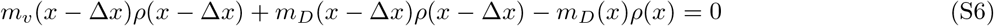

which reduces to

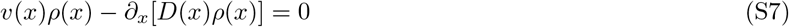

in the continuum limit as expected. Similarly, we assume that there is no influx or outflux of bacteria from diffusion at the right boundary (*x = L*). This yields

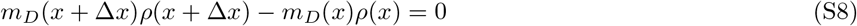

or

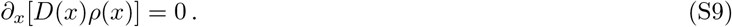

A related set of boundary conditions can be derived by assuming that λ(*x*) is positive only in some finite region of space, and enforcing the physical constraint that *ρ*(*x*) → 0 as *x* → –∞ and *ρ*(*x*) → const as *x* → ∞. We will find that this unbounded formulation can sometimes be more convenient in our later derivations, while yielding qualitatively similar results as the bounded model above.

At steady state, these boundary conditions imply that total growth rate across the colon must equal the outflow at *x = L*,

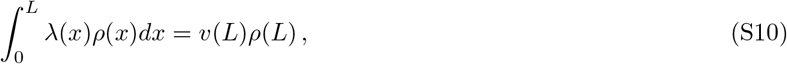

such that the average per capita growth rate is given by

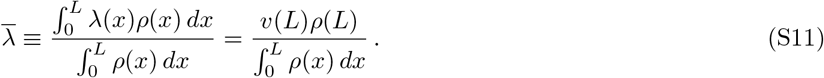

In the limit of a long colon, the resident population will often be dominated by regions with saturated growth 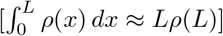, so that

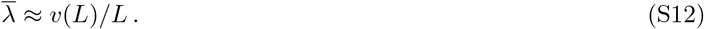

This allows us to recover the standard relationship between the average replication time and the total transit time across the colon (59). We note that all of these calculations assume that the daily elimination of feces does not substantially distrurb the steady-state assumption in Eq. (S5). This is partially justified by estimates that only ~20% of the colonic volume is excreted each day (28). However, further work is required to assess the impact of these daily fluctuations on the steady-state assumptions employed here.

#### Stochastic dynamics of a rare lineage

We are now in a position to derive the stochastic dynamics of a rare mutant lineage when the larger population has reached equilibrium. As long as the mutant is sufficiently rare, we expect it to have a negligible influence on the environmental parameters, *r_i_*(*x,t*), so that λ(*x*) will remain close to its equilibrium form. We will now consider scenarios where the mutant has a of modified birth and death rate, *b_m_* (*x*) and *d_m_*(*x*), which will also be constant in time when the mutant is sufficiently rare. Since the mutant lineage will start out with a small number of cells, its density profile *ρ_m_*(*x,t*) must now be described by a stochastic update rule

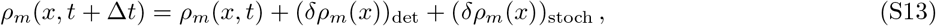

where the deterministic term is similar to the mean-field dynamics in Eq. (S4) above:

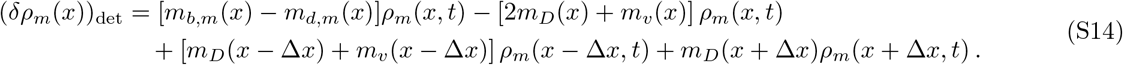

The stochastic term is given by

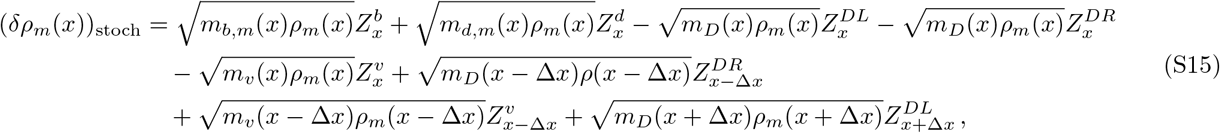

where the 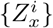 are independent Gaussian random variables with zero mean and unit variance:

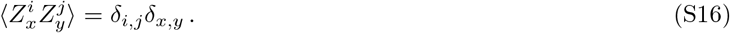

Note that due the advection and diffusion terms, the stochastic term is not diagonal in the *ρ_m_* (*x*) basis, since it will generally include correlations from neighboring *x* values. In our analysis below, we will often be interested in the moments of quantities like ∫*ϕ*(*x*)*δρ_m_*(*x*) *dx*, where *φ*(*x*) is an arbitrary deterministic function. In this case, it is often helpful to diagonalize these integrals in the *ρ_m_*(*x*) basis by rewriting them in the form

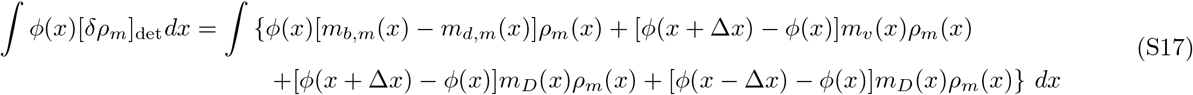

and

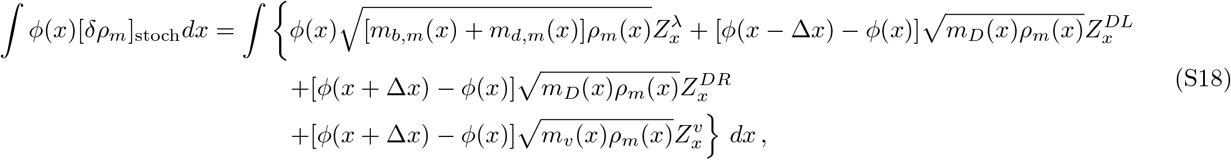

These identities are particularly useful for calculating the generating functional of the mutant lineage, starting from an initial density profile *ρ_m_*(*x*, 0):

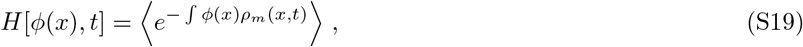

Recursion relations for *H*[*ϕ*(*x*),*t*] can be obtained using standard approaches (57, 58). For example, by considering the change in *H*[*ϕ*(*x*),*t*] over a single timestep Δ*t*, we can rewrite the generating function as

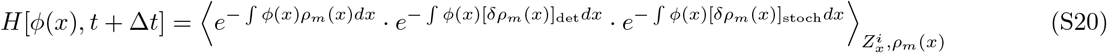

where the angle brackets denote an average over both 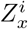 and *ρ_m_*(*x*). By expanding the exponentials to leading order in *m_i_* (*x*) and performing the relevant averages, we obtain a standard system of equations

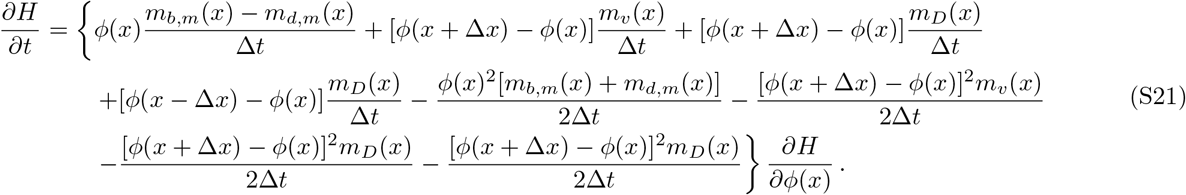

In the continuum limit, this system reduces to the partial differential equation,

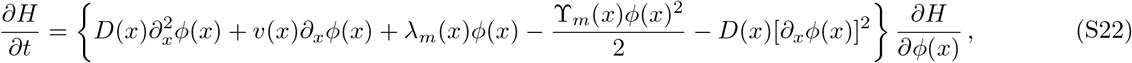

subject to the initial condition

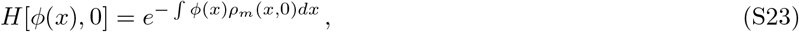

where we have defined the net growth rate,

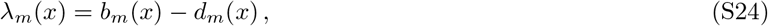

and total variance in offspring number,

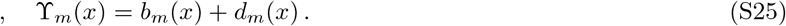

The generating function in Eq. (S22) is identical to the one produced by the Langevin model in Eq. (3). This shows that Eq. (3) can be viewed as the continuum limit of the microscopic model in Eqs. (S14) and (S15). Both models are similar to the linear branching processes that have been analyzed in previous work on evolutionary traveling waves (56, 57, 58). The main difference in this case is the additional nonlinear derivative of *ϕ*(*x*) in Eq. (S22) which arises from the conservative nature of the diffusive fluctuations in space.

The linearity of the noise term in Eqs. (3) and (S22) yields a straightforward equation for the average mutant density profile,

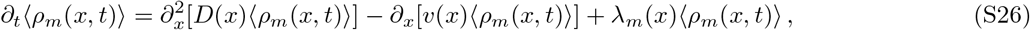

which is similar to the mean field dynamics in Eq. (S4). Formal solutions for the entire distribution of *ρ_m_*(*x,t*) can also be obtained from Eq. (S22) using the method of characteristics. If we let *τ* measure time before the present, then the characteristic curves *ϕ*(*x, τ*) are defined by

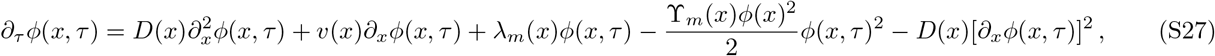

with the initial condition *ϕ*(*x*, 0) = *ϕ*(*x*). The time-dependent generating function is then given by

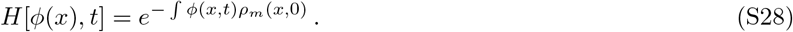

We will explore the behavior of this solution in several specific cases below.

### 2. DYNAMICS OF A NEUTRAL LINEAGE

We first consider the dynamics of a neutral lineage 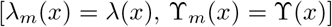, where the characteristic function reduces to

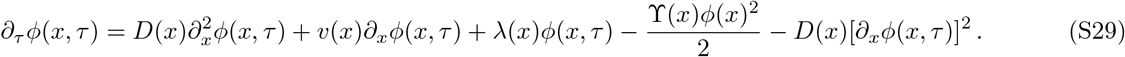

To analyze this equation, it will be useful to first consider the behavior of the linear version,

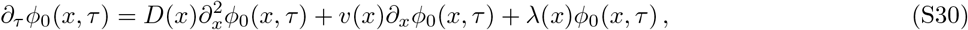

which is obtained by dropping the *ϕ*^2^ and (*∂_x_ϕ*)^2^ terms in Eq. (S29). We note that the right-hand side of Eq. (S29) is the adjoint of the deterministic dynamics for 〈*ρ_m_*(*x,t*)〉 in Eq. (S26). Thus, if *ϕ*_0_(*x, τ*) is a solution of Eq. (S30), then

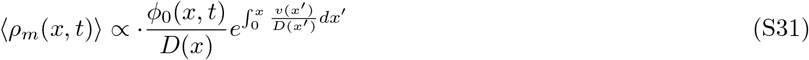

is also a solution for 〈*ρ_m_*(*x,t*)〉 for the particular initial condition

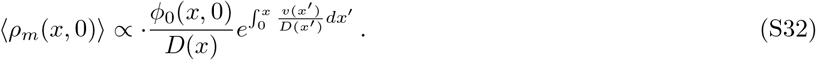

Since the average density profile of a neutral lineage must eventually approach 〈*ρ_m_*(*x, t*)〉 *x ρ*(*x*) at long times, Eq. (S31) shows that *ϕ*_0_(*x, τ*) must also approach a steady state shape as *τ* → ∞. The linearity of Eq. (S30) implies that these steady-state solutions can be expressed in the general form

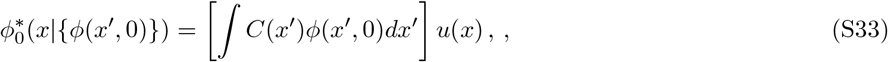

where *C* (*x*) is a weighting function that we determine below, and

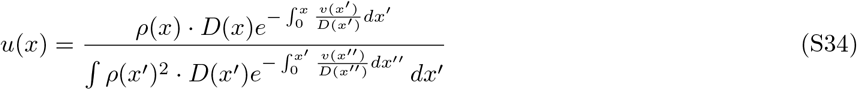

is a solution to the fixed point equation,

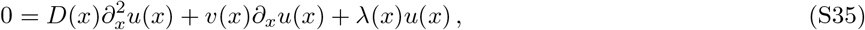

with an overall normalization set by *∫u*(*x*)*ρ*(*x*)*dx* = 1. We have adopted this normalization convention to be consistent with the u(x) function defined in Ref. (37). In that work, the *u*(*x*) function was initially derived in terms of the Greens function of the deterministic dynamics in Eq. (S4). Here we see that it also arises as a solution to the linearized version of the neutral characteristic function in Eq. (S29). We discuss this connection in more detail below.

The linearized solutions in Eq. (S33) will be important because they capture the leading order behavior of the characteristic curves *ϕ*(*x, τ*) in the limit of small *ϕ*(*x*, 0). For example, by substituting Eq. (S33) into Eq. (S28), we find that the leading order contribution to the generating function is given by

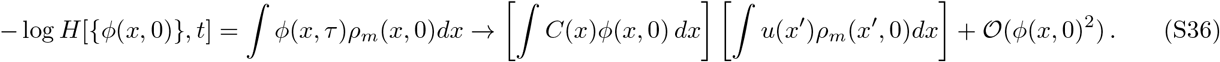

The average mutant density profile, 〈*ρ_m_*(*x,t*)〉, then follows as

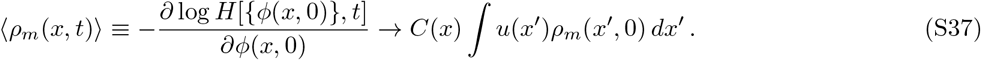

In particular, this shows that if *ρ_m_*(*x*, 0) is proportional to the steady state density profile, *ρ_m_*(*x*, 0) = *fρ*(*x*), then 〈*ρ_m_*(*x,t*)〉 must be constant over time:

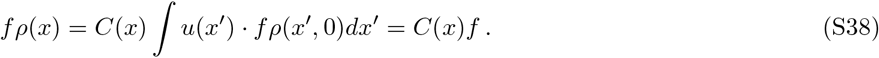

This implies that *C*(*x*) = *ρ*(*x*), and hence

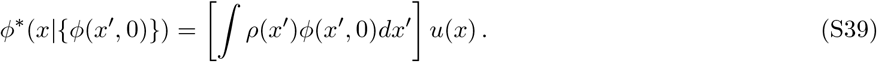

The functional form of Eq. (S39) implies that *ρ*(*x*’)*u*(*x*) is the Greens function for the linearized dynamics in Eq. (S30) in the limit that *τ* → ∞, just as *ρ*(*x*)*u*(*x*’) is the Greens function for the deterministic dynamics of 〈*ρ_m_*(*x, t*)〉 (37).

We will let *τ*_mix_ denote the “mixing timescale” required for *ϕ*_0_(*x, τ*) to approach its fixed point *ϕ*\*(*x*). This timescale can have a complicated dependence on λ(*x*), *v*(*x*), and *D*(*x*), but due to the linearity of Eq. (S30), it must be independent of the overall scale of *ϕ*(*x*, 0) – or equivalently, of *ρ_m_*(*x,t*). This suggests that in sufficiently large populations, we will be able to exploit a separation of timescales between the linearized growth dynamics (*τ*_mix_) and the slower evolutionary process of genetic drift [∝ 1/ *∫ρ_m_*(*x,t*)*dx*].

#### 1. Variance accumulated due to genetic drift

As an example of this separation-of-timescales approach, we note that the variance of *ρ_m_*(*x, t*) can be obtained from the second order corrections to *ϕ*(*x, τ*) in the limit that *ϕ*(*x*, 0) is small:

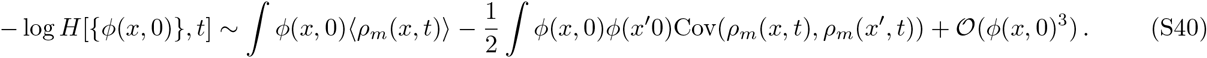

We can calculate these contributions by computing the next order terms in the perturbation expansion for *ϕ*(*x, τ*) above. By treating the nonlinear terms as a small correction, a basic iterative solution shows that the *τ* ≫ *τ*_mix_ behavior is given by

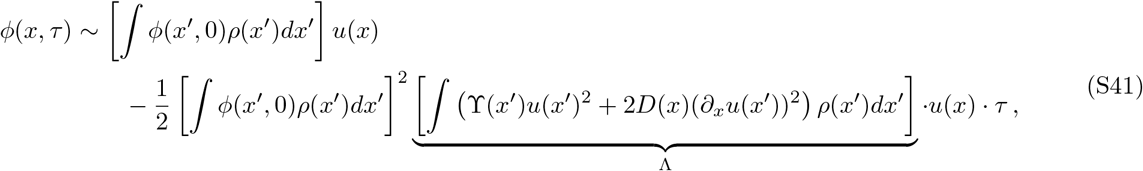

where Λ coincides with the effective rate of genetic drift in Eq. (7) in the main text. This iterative solution in Eq. (S41) is self-consistently valid when

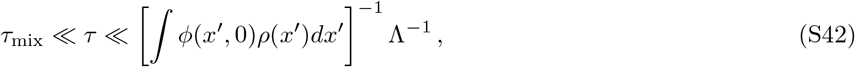

which can be satisfied for any fixed value of *τ* ≫ *τ*_mix_ by choosing sufficiently small values of *ϕ*(*x*, 0). Note, however, that the converse is not true: for any fixed value of *ϕ*(*x*, 0) ≪ 1/Λ*τ*_mix_, the perturbative solution in Eq. (S41) will eventually break down for sufficiently large *τ*.

Fortunately, the variance of *ρ_m_*(*x,t*) can be extracted from the generating function *H*[{*ϕ*(*x*)},*t*] by choosing a sufficiently low value of *ϕ*(*x*, 0) = *ϕ*(*x*) so that Eq. (S41) remains valid for each desired value of *t*. By substituting this solution into Eq. (S28), we see that the second order contributions to the generating function are given by

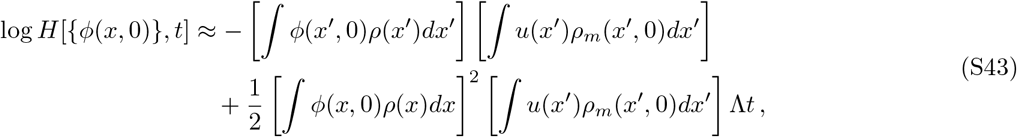

so that

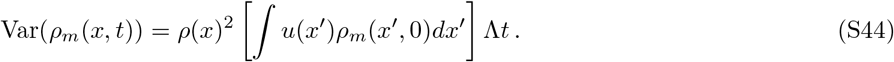

This solution is self-consistently valid as long as *t* ≫ *τ*_mix_. By expressing this result as a coefficient of variation:

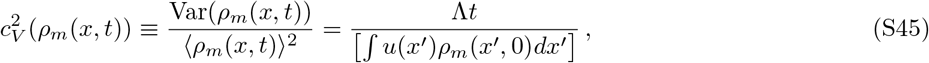

we obtain a natural definition of the drift timescale,

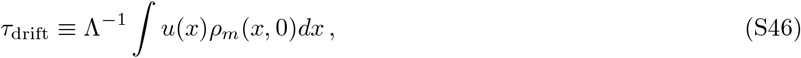

based on the time that the coefficient of variation first exceeds one.

#### 2. Survival probability at long times

It will also be useful to calculate *ϕ*(*x,τ*) on timescales much longer than *τ*_drif_ to This is particularly important for calculating the survival probability of a lineage at long times. This survival probability can be calculated from an asymptotic expansion of the moment generating function in the limit that *t* ≫ *τ*_drift_ *τ*_mix_:

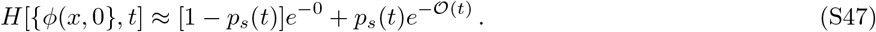

Plugging in our method-of-characteristics solution for *H*[*ϕ*(*x*),*t*] above, we find that

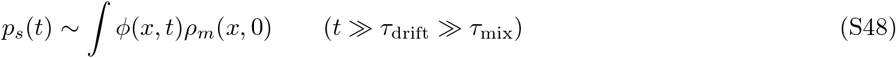

In this case, we can no longer calculate *ϕ*(*x, τ*) by appealing to the simple perturbative expansion above, since the nonlinear terms (which represent the effects of fluctuations) will eventually become important for sufficiently large *τ*. Nevertheless, due to the separation of timescales, we expect that *ϕ*(*x,t*) will remain close to *ϕ*\*(*x*) in a sense that we will make more precise below. At long times, we know that *ϕ*(*x, t*) must eventually vanish (since neutral mutations must always go extinct in an infinitely large population). Motivated by the behavior in the single locus case (54, 55), we define a new function *ψ*(*x,τ*) such that

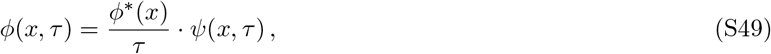

anticipating that *ψ*(*x,τ*) approach some time-independent shape in the limit of large *τ*. In fact, it will be useful to split up *ψ*(*x,τ*) into three independent components:

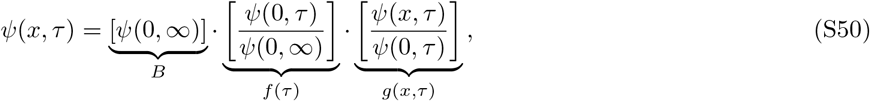

where *B* is an arbitrary constant and *f* (*τ*) and *g*(*x, τ*) are unknown functions that satisfy the boundary conditions

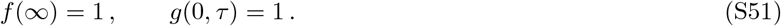

The spatial boundary conditions on *ϕ*(*x, τ*) provide an additional set of boundary conditions for the function *g*(*x, τ*). At *x* = 0, we have

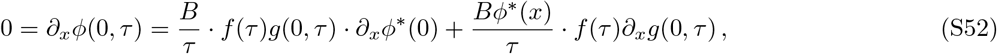

which implies that

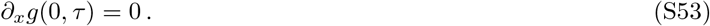

Likewise, at *x* → ∞ we must have *ϕ*(*x, τ*) → 0, which implies that

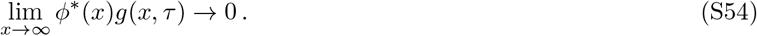

Substituting these definitions into Eq. (S29), we obtain a corresponding characteristic curve for *g*(*x, τ*):

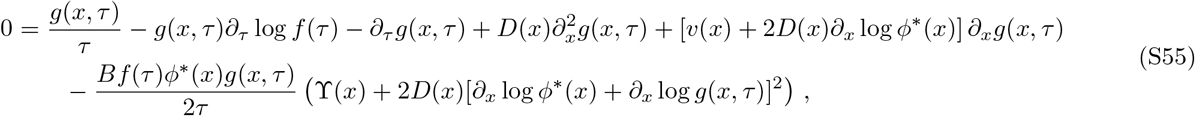

subject to the boundary conditions

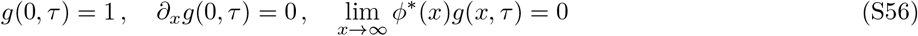

We now seek an asymptotic solution of Eq. (S55) in the limit of large *τ*. We assume that *f* (*τ*) and *g*(*τ*) can be written as a power series in 1/*τ*:

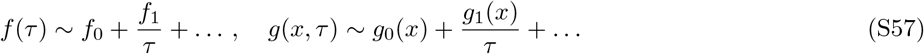

The boundary conditions on *f* (*τ*) and *g*(*x, τ*) imply that

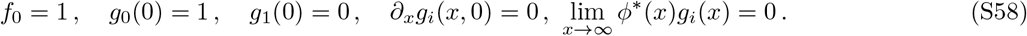

After substituting these expansions into Eq. (S55) and collecting like powers of 1/*τ*, we find that the zeroth order contribution yields:

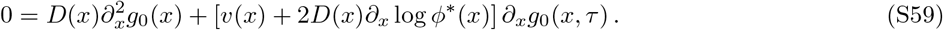

The only solution that satisfies the boundary conditions at *x* = 0 and *x* = ∞ is

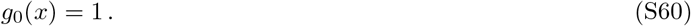

The next order contribution from Eq. (S55) then yields

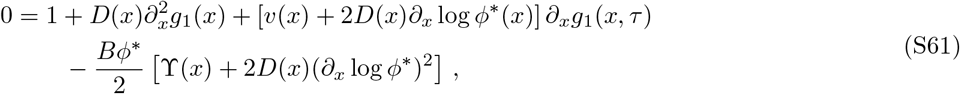

or

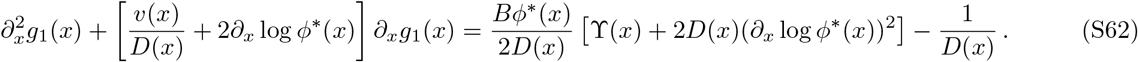

This is a first order equation for *∂_x_g*_1_(*x*,), which has the solution

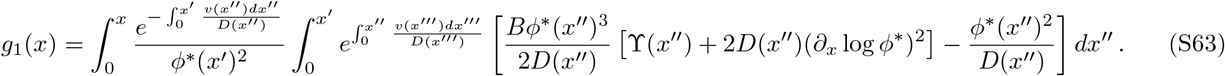

The limits of integration in Eq. (S63) were chosen to satisfy the boundary conditions *g*_1_(0) = 0 and *∂_x_g*_1_(0) = 0.

To apply the remaining boundary condition at *x* = ∞, it is helpful to impose the mild assumption that λ(*x*) eventually vanishes for sufficiently large *x*. This implies that the solutions for *ϕ*\*(*x*) must scale as

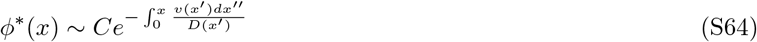

for sufficiently large *x*. This means that the prefactor inside of the first integral must correspondingly grow as 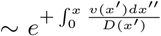. Similarly, at large *x*, the inner integral must eventually approach a constant value at large *x*. The only way that the combined expression can grow more slowly than 1/*ϕ*(*x*) is if the inner integral vanishes at *x* → ∞:

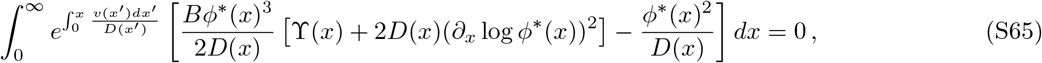

which requires that

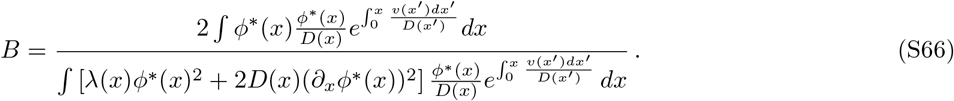

Plugging this back into our expression for *ϕ*(*x,τ*), and recalling that

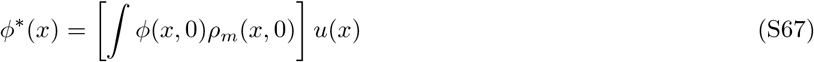

and

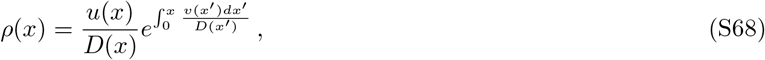

we find that the characteristic curve *ϕ*(*x, τ*) reduces to

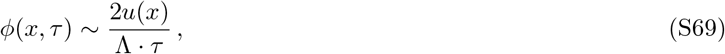

where Λ is the effective rate of genetic drift defined in Eq. (7). The survival probability is therefore given by

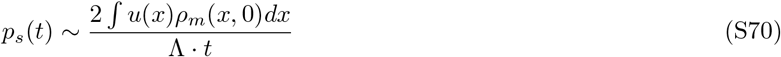

We can combine this result with the average lineage size above to obtain the average size conditioned on survival:

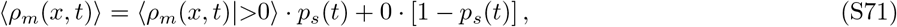

which yields

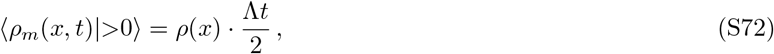

or

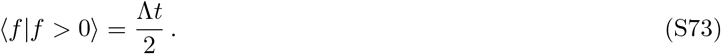

Note that the dependence on the initial density profile *ρ_m_*(*x*, 0) cancels out, so that the conditional mean is independent of the initial conditions for times *t* ≫ *τ*_drift_ ≫ *τ*_mix_. Similarly, the conditional variance is given by

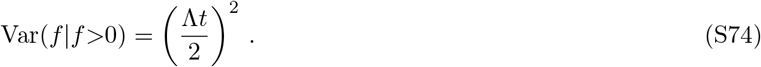

Motivated by these results, we can also solve for the full distribution of *f* (*t*) conditioned on survival. The key insight will be to restrict our evaluation of the generating function regimes where

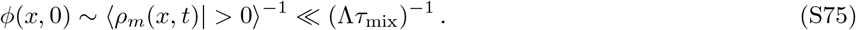

Equations (S73) and (S74) show that this will be sufficient to capture the typical values of *ρ_m_*(*x,t*), though it will break down for smaller sizes where the lineage is close to extinction. By restricting our attention to this regime, we can again employ the short-time expansion that we used to derive the mean and variance in above. At intermediate times *τ*_mix_ ≪ *τ* ≪ Λ^-1^, the generating function will quickly reduce to

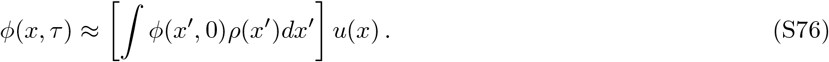

Thus, it will be sufficient to consider effective initial conditions of the form *ϕ*(*x*, 0^+^) = *z*_eff_*u*(*x*). The next order correction yields

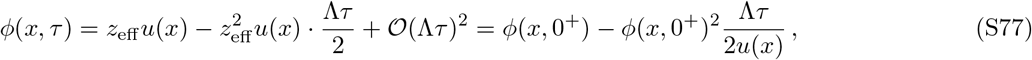

which can be recast as a coarse-grained differential equation for *ϕ*(*x,t*):

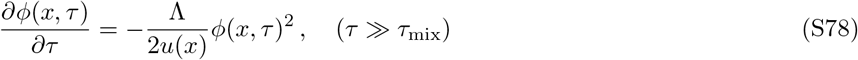

After solving this equation, and applying the initial condition 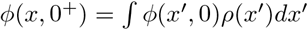, we find that

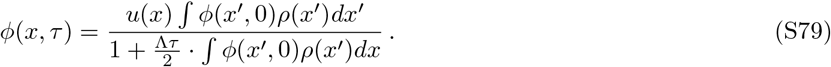

The generating function for the total mutant frequency can be obtained by setting *ϕ*(*x*, 0) = *z/N*. This yields

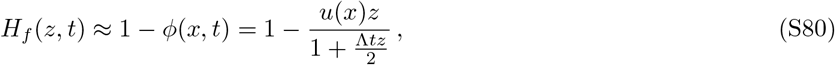

and a corresponding survival probability

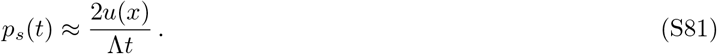

The conditional generating function is therefore given by

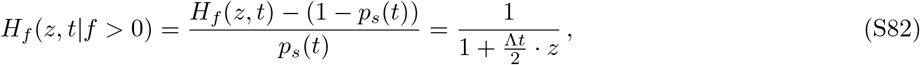

which is an exponential distribution with mean Λ*t*/2. These expressions are equivalent to a well-mixed Wright-Fisher model,

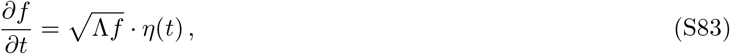

with an effective initial frequency *f* (0) = *u*(*x*) and an effective rate of genetic drift Λ.

#### 3. Effective generation time

We can use this result to calculate the effective generation time by focusing on the long-term substitution rate of neutral mutations. In principle, mutations can arise through a mixture of replication errors during cell division and DNA damage between divisions. We can encapsulate both of these processes within our model using a spatially varying mutation rate,

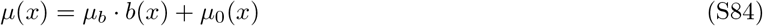

where *b*(*x*) is the local birth rate, *μ_b_* is the mutation rate from cell division, and *μ*_0_(*x*) is the background mutation rate from DNA damage. Once a mutation occurs, our previous analysis shows that it will survive to times *t* ≫ *τ*_mix_ with probability

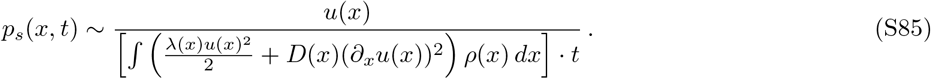

At this point, the mutant and wildtype populations will have equivalent spatial distributions (since *t* ≫ *τ*_mix_).

Symmetry considerations then imply the ultimate fixation probability of the neutral mutant must be proportional to its relative size, so that

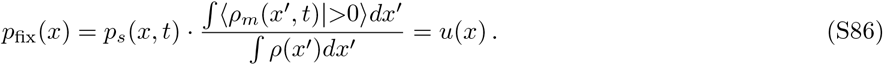

This allows us to identify *u*(*x*) with the fixation probability of a neutral lineage, as previously noted by Ref. (37). The long-term substitution rate of these neutral mutations is therefore given by

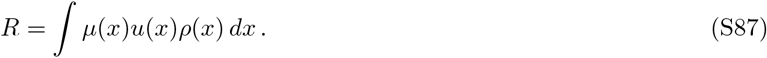

We can connect this to the effective generation time by focusing on the subset of division-marking mutations, for which *μ*(*x*) = *μ_b_* · *b*(*x*), and

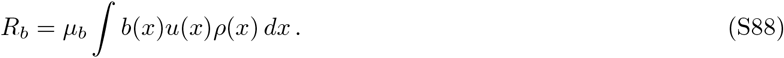

By definition, *R_b_* must also be equal to *μ* · *t/τ_g_*, where *τ_g_* is the effective generation time. This yields an expression for *τ_g_* in terms of an average over *u*(*x*) and *ρ*(*x*),

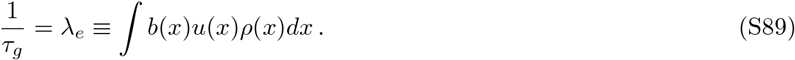

Equation (9) in the main text follows from this expression by replacing the local birth rate with its equivalent definition, *b*(*x*) = λ(*x*) + *d*(*x*). We can use this result to re-express the rates of genetic drift above in more traditional units of generations:

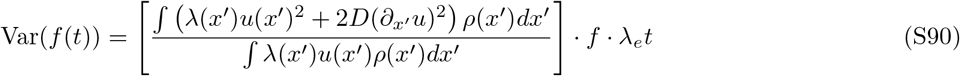

The quantity in braces can then be identified as an “effective population size”

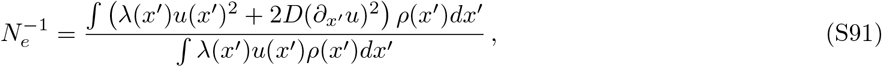

which is equivalent to Eq. (10) in the main text. These definitions allow us to rewrite our earlier results in a particularly compact form:

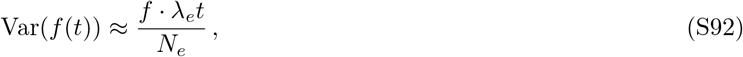

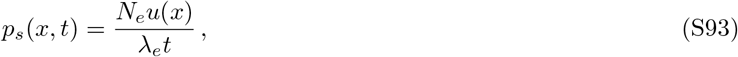

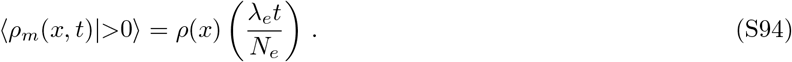

Together, these results show that on timescales much longer than *τ*_mix_, the dynamics of neutral lineages resemble a Wright-Fisher model with effective generation time 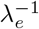 and effective population size *N_e_*.

### 3. DYNAMICS OF A SELECTED LINEAGE

We can use a similar set of approaches to characterize the dynamics of selected mutations. In this case, the characteristic curve in the generating function is now given by

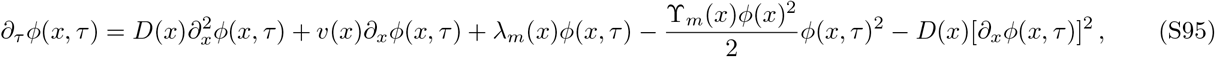

where λ_*m*_(*x*) is the position-dependent growth rate of the mutant strain. If λ_*m*_ (*x*) is sufficiently close to the wildtype growth rate λ(*x*), then the dynamics of *ϕ*(*x, t*) on timescales 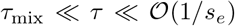 can be derived using the same short-time perturbation approach as above. The first order contribution is given by

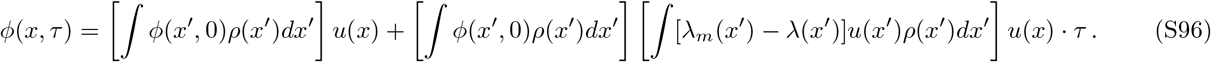

The average mutant density profile is therefore given by

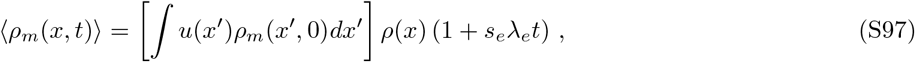

where

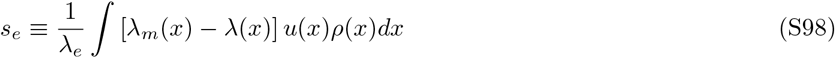

is the effective selection coefficient defined in Eq. (12) in the main text. Iterating over multiple short time windows then yields

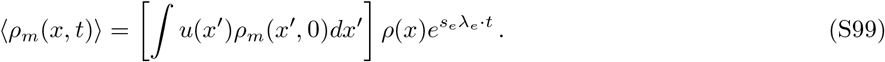

For a constant growth rate advantage, λ_*m*_(*x*) = (1 + *s*)λ(*x*), we find that

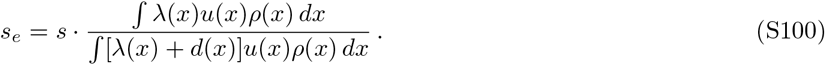

This shows that the effective selection strength is proportional to s as expected. When cell death is negligible in the regions that dominate λ(*x*)*u*(*x*)*ρ*(*x*), this further reduces to *s_e_ = s*.

#### 1. Establishment probability

In contrast to the neutral scenario above, the characteristic curve for a beneficial mutation must eventually saturate at a nonzero fixed point:

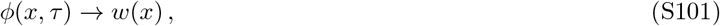

described by the ordinary differential equation in Eq. (13). For small growth rate differences, we can solve for the fixed point *w*(*x*) using a perturbation expansion that is structurally similar to the one we used in the neutral case above. We first introduce a perturbation scale e such that

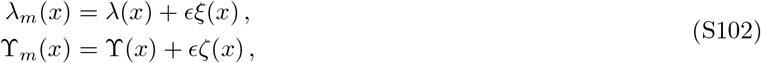

with *ϵ* ≪ 1. We then rewrite *w*(*x*) as

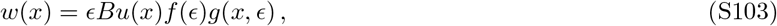

where *u*(*x*) is the linearized neutral fixed point above, and *f* (*ϵ*) and *g*(*x, ϵ*) are unknown functions that satisfy

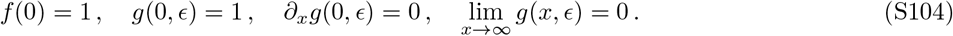

Substituting these expressions into Eq. (13) in the main text, we obtain an analogous differential equation for *g*(*x, ϵ*):

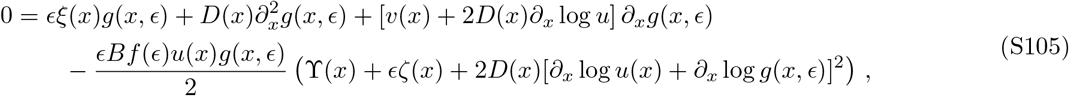

subject to the boundary conditions

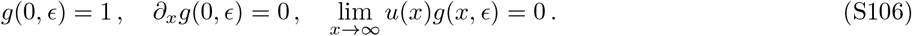

We then assume an analogous set of asymptotic expansions:

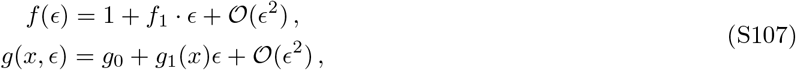

and substitute these expressions into Eq. (S105). At zeroth order in *ϵ*, we find that

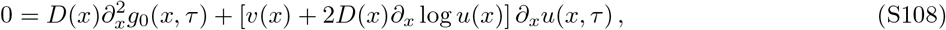

and hence *g*_0_ = 1 as above. At first order in *ϵ*, we have

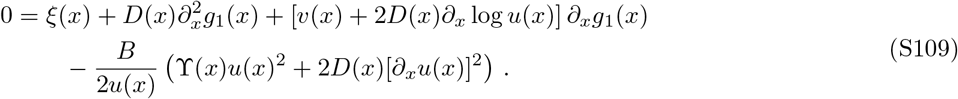

This is a similar first order equation as in the neutral case, and the solution is given by

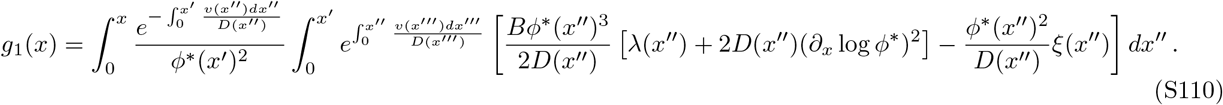

As above, the finite boundary condition at *x* = ∞ requires that B is chosen to satisfy

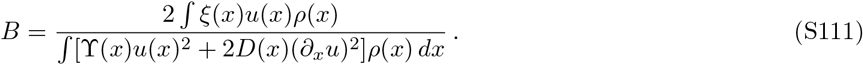

This implies that

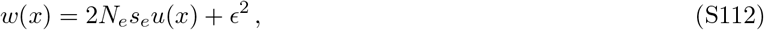

as expected from Eq. (11).

We can test the limits of this perturbation expansion by turning to a numerical solution of Eq. (13). In this case, it is helpful to work in logarithmic space by defining

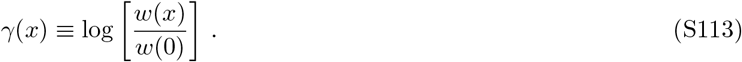

This function satisfies the related differential equation

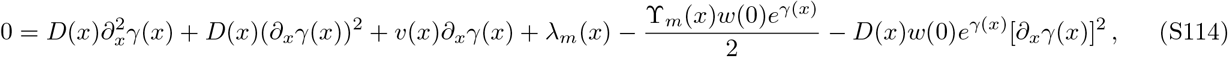

with the initial conditions

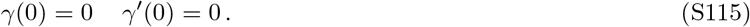

We will often be interested in scenarios where there is a critical position *x* = *x*_0_ above which λ_*m*_(*x*) and 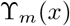 vanish, and where *v*(*x*) and *D*(*x*) eventually attain constant values, *v*(*x*) = *v* and *D*(*x*) = *D*. When this occurs, the solution to Eq. (S114) for *x > t* is given by

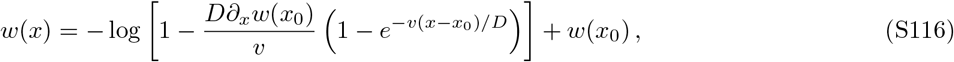

where the initial conditions *w*(*x*_0_) and *∂_x_w*(*x*_0_) must be chosen to ensure that *w*(*x*) → 0 as *x* → ∞. By examining the form of Eq. (S116), we see that the only way that this can happen is if *w*(*x*_0_) and *∂_x_w*(*x*_0_) are related by

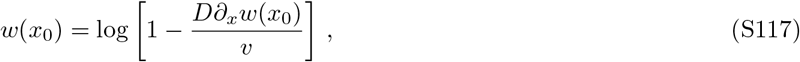

which implies that

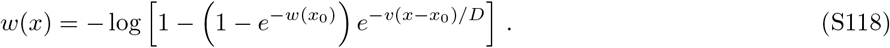

In terms of the logarithm *γ*(*x*), these expressions become

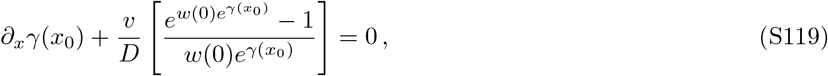

and

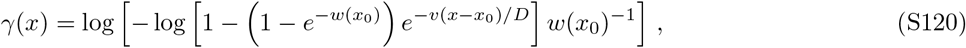

respectively. These expressions allow us to obtain numerical solutions for the fixation profile by first solving for *w*(*x*) over the finite region 0 ≤ *x* ≤ *x*_0_ and then enforcing continuity with this analytical solution for the large-*x* limit at *x* = *x*_0_. We used this approach to calculate the large-*s_e_* behavior of the fixation probability in Figs 3 and 4, which is described in more detail in Supplementary Information 4.4 and 5 below.

#### 2. Long-term growth rate of strongly beneficial mutations

We can use a related approach to investigate the large-*s_e_* behavior of the average mutant density profile in Eq. (S26). We assume that the long-time behavior can be written in the form

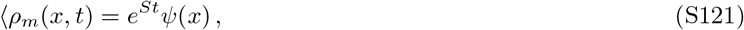

where *S* is an effective exponential growth rate and *ψ*(*x*) is an arbitrary spatial profile. Substituting this ansatz into Eq. (S26), we find that *ψ*(*x*) must satisfy the related differential equation

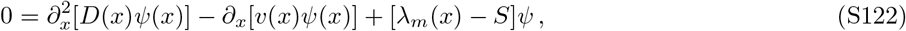

and is subject to the same boundary conditions as 〈*ρ_m_*(*x,t*)〉:

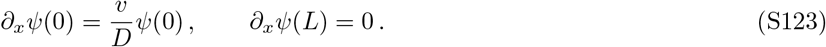

When λ_*m*_(*x*) is close to λ(*x*), we can use a perturbative solution of Eq. (S122) to obtain an alternate derivation of the effective selection coefficient in Eq. (12) in the main text.

To do so, we first note that the duality between *ρ*(*x*) and *u*(*x*) holds for *ψ*(*x*) as well. This motivates us to define a function

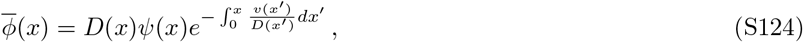

which satisfies the dual equation

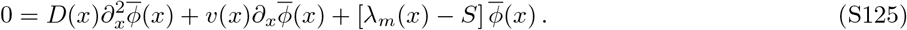

This is similar to Eq. (13) in the main text, except with 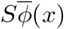 replacing the 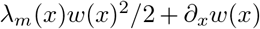 term. Thus, by repeating our perturbative calculation for *w*(*x*) above, we conclude that *S* must be chosen to satisfy

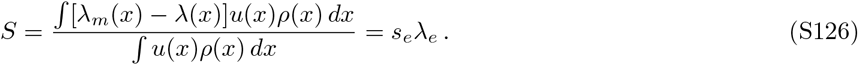

in agreement with the separation-of-timescales result in Eq. (12).

The advantage of Eq. (S122) is that it can still be used to solve for the exponential growth rate even when *Sτ*_mix_ ≫ 1.

We carry out this calculation for the minimal model in Fig. 3 in Supplementary Information 4.5 below.

### 4. ANALYTICAL SOLUTIONS FOR A MINIMAL MODEL OF BACTERIAL GROWTH

We can gain additional intuition for our results by considering a concrete model of microbial growth in the gut, where many of the important quantities can be determined analytically. In this simplified model, we assume that the flow velocity and mixing rates are spatially uniform, so that

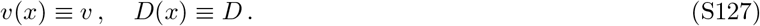

We also assume that the gut is long enough that the total length *L* can be regarded as effectively infinite (*vL/D* → ∞). We neglect cell death, and assume that microbial growth depends on a single nutrient that is supplied at the entrance of the colon, which is consumed by cells during division. We assume that the growth rate has a step-like dependence on the local nutrient concentration,

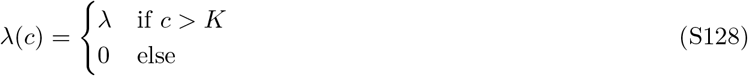

with a maximum growth rate λ and saturation constant *K*. This function is qualitatively similar to the Monod-like models that have been employed in previous work (28, 65), but with a sharper transition between the no-growth and saturated-growth-rate regimes. This will turn out to be a crucial simplification that will allow us to obtain analytical solutions for the growth rate and density profiles that are attained at steady state.

The assumptions of our model imply that the local nutrient concentration must be a monotonically decreasing function of *x*, since the nutrients are progressively consumed as they flow down the colon. The step-like growth function then implies that the local growth profile λ(*x*) must also develop a step-like form,

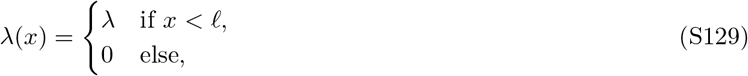

which transitions from high-growth to no-growth at a critical position *x = ℓ*. The location of this transition is determined by the point where the local nutrient concentration first drops below *c*(*x*) = *K*. This depends in turn on the local density profile *ρ*(*x*), which dictates the total consumption of resources between *x* = 0 and *x* = *ℓ*.

An important feature of our minimal model is that it allows us to side-step this coupled calculation of *n*(*x*) and *ρ*(*x*). To see this, we note that the solutions for the steady-state density profile in Eq. (S5) will have two degrees of freedom, one of which is fixed by the overall normalization condition,

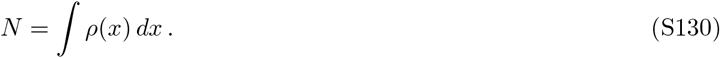

Since there are two independent boundary conditions for *ρ*(*x*) – and only one remaining degree of freedom — we see that *ℓ* can be uniquely determined by the requirement that Eq. (S5) admits a non-trivial solution. Furthermore, the solution to this “eigenvalue problem” also determines the shape of the density profile *ρ*(*x*) up to an overall normalization factor, which is fixed by Eq. (S130). This shows that the explicit dependence on the nutrient profile is completely encapsulated by the population size *N*, which will vary as a function of the nutrient input flux and the saturation constant *K*. However, for a fixed value of *N*, the steady-state growth and density profiles are completely determined by Eq. (S5). We adopt this perspective in our calculations below, treating *N* as an independent parameter that must be determined experimentally.

We note that this same argument applies for any model that generates a single parameter family of growth profiles like Eq. (S129). This suggests that Eq. (S129) may be considerably more general than the specific growth model proposed above, and can also be viewed as a purely phenomenological approximation to the empirical growth profiles in Fig. 4.

#### 1. Steady state density profile

To solve for the steady-state density profile, it will be convenient to switch to diffusion units by defining the rescaled variables:

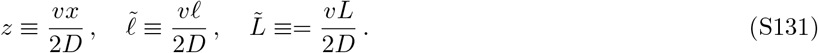

Equation (S5) can then be written in the dimensionless form

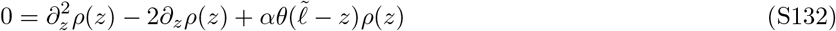

with *α* defined by

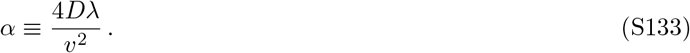

Equation (S132) admits a simple piecewise solution for 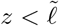 and 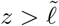, respectively. In the bulk of the colon (*z* > *ℓ*), the density profile satisfies

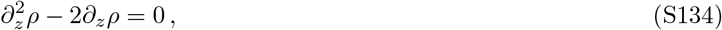

which has the general solution

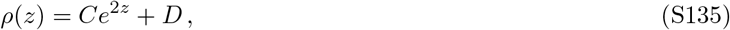

where *C* and *D* are integration constants. The finite boundary condition at 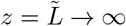 requires that *C* = 0, so that *ρ*(*z*) is simply a constant in this region.

On the other hand, when 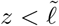, the density profile instead satisfies

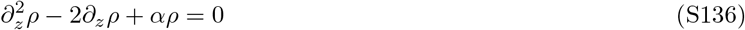

which has the general solution

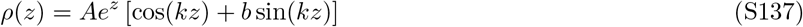

where

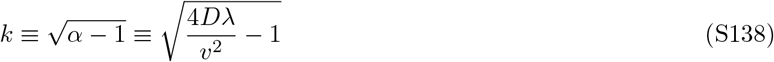

and *A* and *b* are a pair of integration constants. Applying the boundary condition at *z* = 0 yields

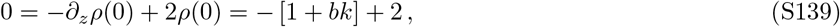

so that *b* = 1/*k* and

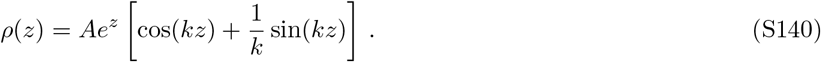

Finally, enforcing continuity at 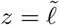 allows us to solve for *D* in terms of *A*:

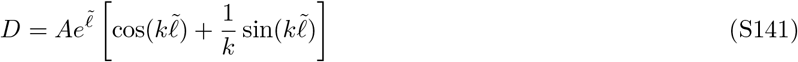

while enforcing continuity of the derivative at 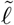 yields an “eigenvalue” equation for 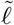:

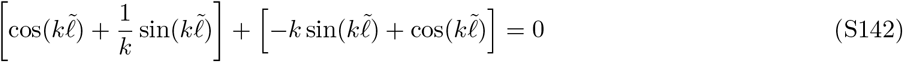

This can be rewritten in the compact form

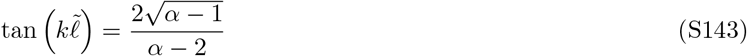

whose solution is given by

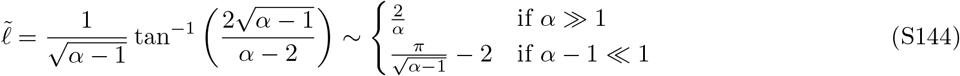

Given this value of 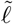, the full piecewise solution for the density profile can be written as

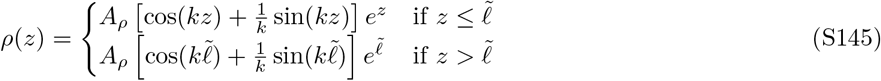

where the normalization constant *A_ρ_* is determined by the overall population *size ∫ ρ*(*z*) *dz* = *N*. Equation (6) then implies that the neutral fixation probability *u*(*x*) is given by

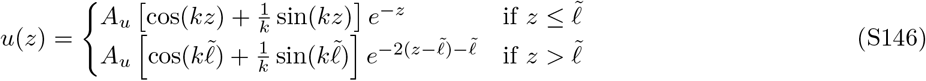

where the normalization constant *A_u_* is fixed by the condition that *∫ u*(*z*)*ρ*(*z*) *dz* = 1. The ancestral distribution *g*(*z*) = *u*(*z*)*ρ*(*z*) then takes on the simple form

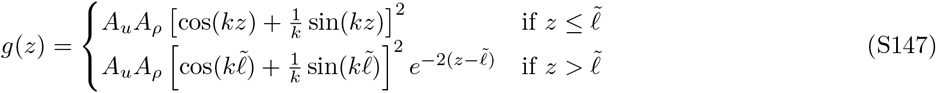

which is independent of *N* as expected and normalized so that *∫ g*(*z*)*dz* = 1.

#### 2. Effective generation time and population size

To calculate the effective generation time and population size, it will be helpful to first obtain explicit expressions for the normalization factors *A_ρ_* and *A_u_* above. In the limit that the length of the colon is very long 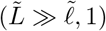, the total population size will be dominated by slowly growing regions 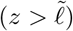, so that

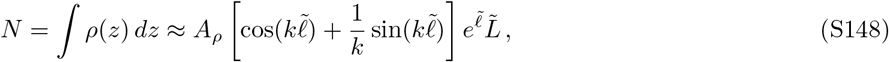

or

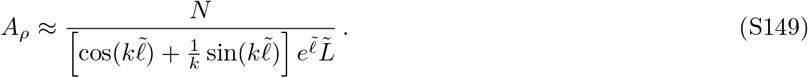

Similarly, the normalization condition *∫ g*(*z*)*dz* = 1 implies that

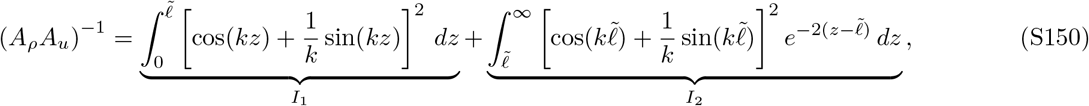

which we have split into contributions from the integrals *I*_1_ and *I*_2_. The latter is given by

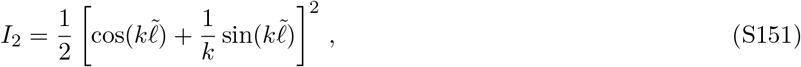

while the former evaluates to

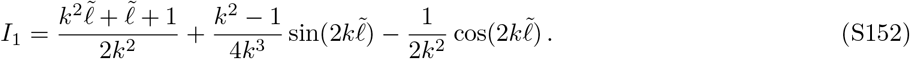

Using these expressions, we can write the effective generation time in the form

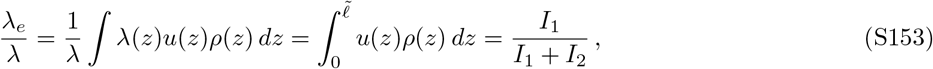

where *I*_1_ and *I*_2_ are defined as above. This yields an exact expression,

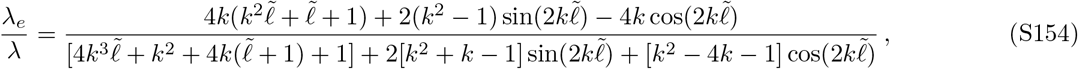

whose large and small *α* limits are given by

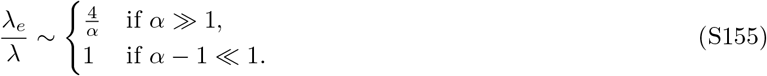

This asymptotic behavior can also be obtained by considering the *I*_1_ and *I*_2_ integrals directly. When *α* ≫ 1 is large, the sine and cosine terms sum to ≈1 throughout the entire interval 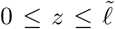. This implies that 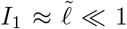 and *I*_2_ ≈ 1/2 and hence

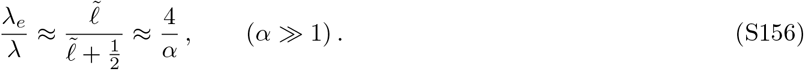

In the opposite extreme (*α* – 1 ≪ 1, *k* ≪ 1, and 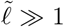), the sine term dominates over the cosine term through the bulk of the interval 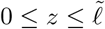, so that

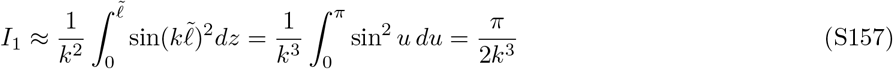

Meanwhile, Eq. (S142) shows that the sine and cosine terms sum to ≈ 1 at 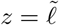, so that 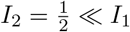, and

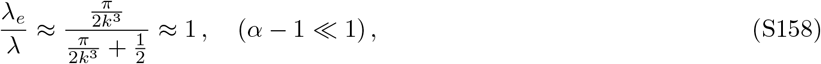

as desired.

We can use a similar strategy to evaluate the effective population size in Eq. (10). We begin by rewriting this expression in the form

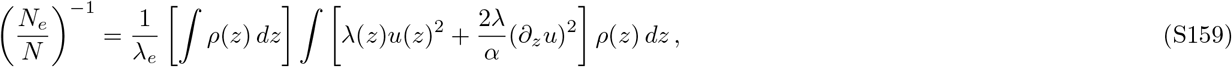

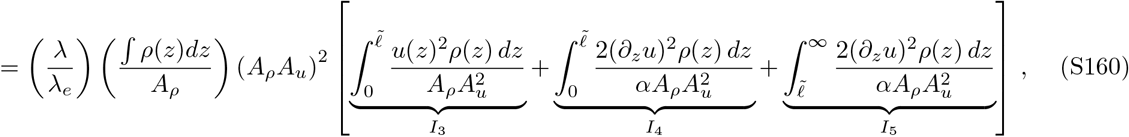

which we have split into contributions from the integrals *I*_3_, *I*_4_, and *I*_5_. To evaluate these expressions, it is helpful to use Eq. (S146) to show that

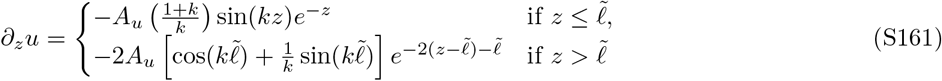

This implies that *I*_5_ is given by

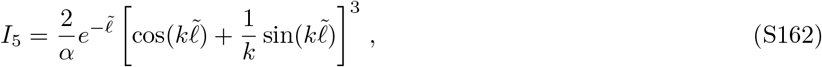

while *I*_3_ evaluates to

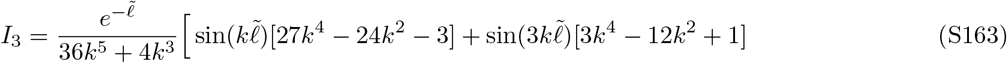

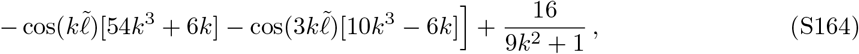

and *I*_4_ is given by

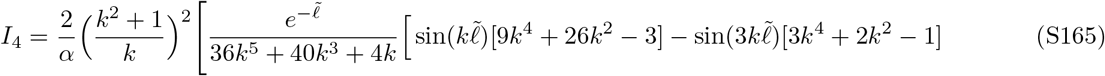

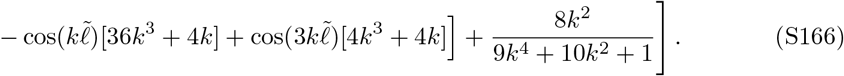

Putting everything together, we have

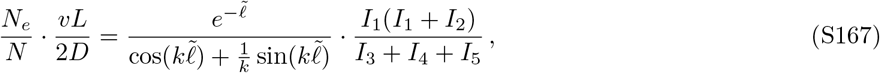

whose large and small *α* limits are given by

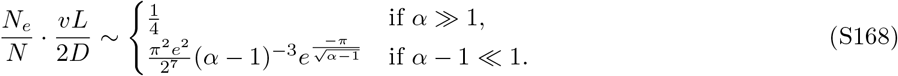

As above, these large and small α limits can also be obtained by considering the individual integrals directly. When *α* ≫ 1 the sine and cosine terms sum to ≈1 (and 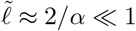), so

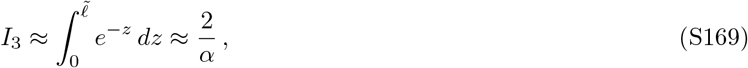

and

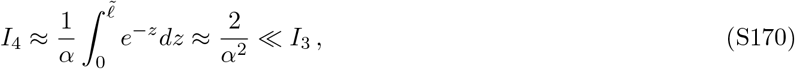

while *I*_5_ evaluates to

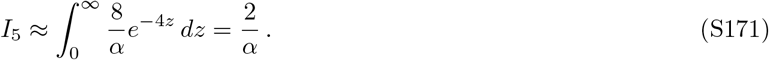

This implies that growth below 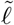 (represented by *I*_3_) and diffusion above 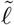 (represented by *I*_5_) make equal contributions to the magnitude of genetic drift. Combining this with our asymptotic expressions for *I*_1_ and *I*_2_ above, we find that

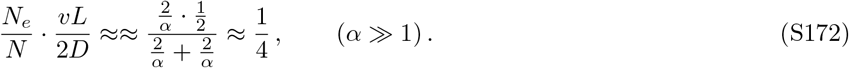

This shows that while the wall-clock rate of genetic drift (Λ) declines for large *α*, the longer generation times in this regime end up compensating for this effect to yield a constant value of *N_e_/N*.

In the opposite extreme (*α* – 1 ≪ 1, *k* ≪ 1, and 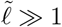), the *I*_3_ integral will be dominated by a narrow region where 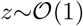, so that

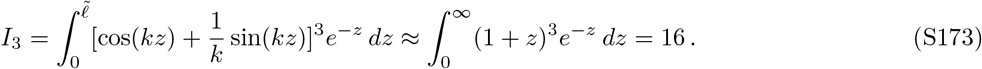

and

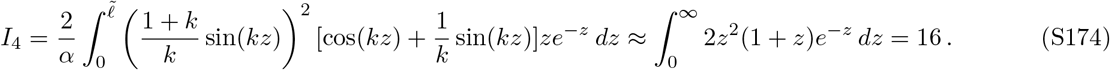

Finally, since 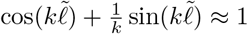, the contribution from *I*_5_ becomes

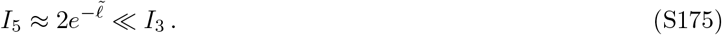

Combined with our expressions for *I*_1_ and *I*_2_ above, this yields,

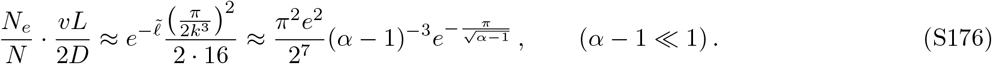

This shows that in the *α* – 1 ≪ 1 regime, genetic drift is dominated by a narrow slice of the ancestral distribution with 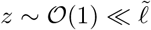. Since *ρ*(*z*) is exponentially increasing in this regime, the bacterial density in this narrow slice of *z* values is also smaller than *ρ*(*t*) by an additional factor of ~ exp(—*ℓ*). This leads to an effective population size that is significantly smaller than the total number of dividing cells, which scales like

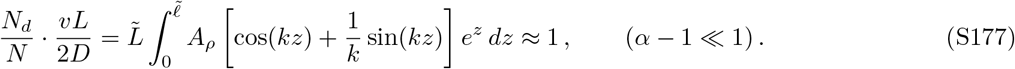

#### 3. Extension to multiple growth regions

To better understand the mechanisms responsible for driving the enhanced rates of genetic drift near the washout threshold (*α* → 1), we considered a generalization of our minimal model that includes a washed out region at the proximal end of the colon. We reasoned that such a model would allow us to determine whether washout itself (*α* < 1) — vs an extended region of near-washout (*α* → 1) growth — is the primary cause of the behavior in Fig. 2F. For simplicity, we assumed that the washed-out region extends all the way from *x* = 0 to *x* = —∞, so that it would have the largest possible impact on the emergent evolutionary parameters. Our generalized model assumes that the growth and flow velocity profiles can be written the piecewise form,

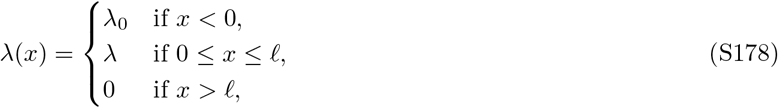

and

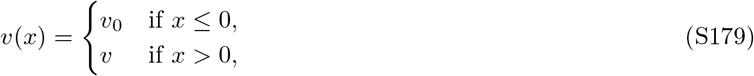

where λ_0_ and *v*_0_ represent the growth and flow parameters in the washed out region.

To solve for the steady-state population density profile, it is helpful to adopt the same units as in Eq. (S131) above, where all lengths are measured relative to the mixing lengthscale in the original the *x* > 0 region. The equation for the steady-state density profile in the new region can then be written in the dimensionless form

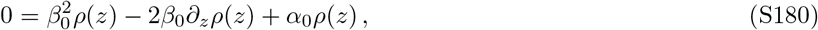

with *α*_0_ and *β*_0_ defined by

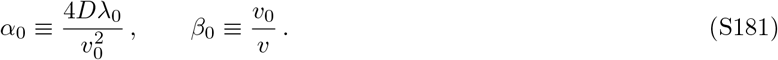

Here, we will be primarily interested in parameter combinations that prevent the population from fully establishing in the *x* < 0 region (*α*_0_ < 1) and where the flow velocity is at least as large as in the *x* > 0 region (*v*_0_ > *v*).

The general solution to Eq. (S180) is given by

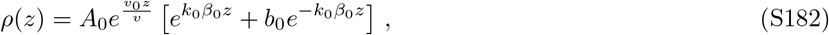

where *A*_0_ and *b*_0_ are integration constants, and

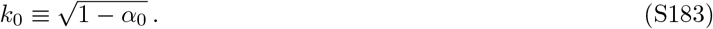

The reflecting boundary condition at *z* = —∞ requires that *b* = 0, so that

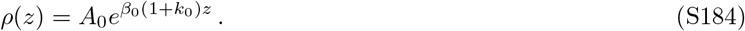

By matching this solution to our previous solution for *ρ*(*z*) in the *z* > 0 region,

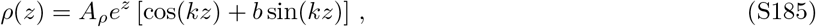

we find that

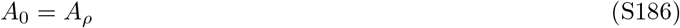

The derivative condition is slightly more complicated in this case, since *v*(*x*) changes abruptly at *x* = 0. In this case, the conservation of particle flux across the boundary yields a generalized continuity condition,

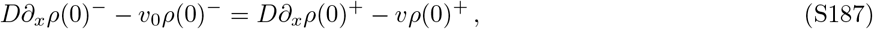

which, in dimensionless form, becomes

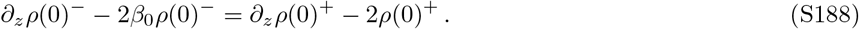

By applying this condition to Eqs. (S184) and (S185), we find that

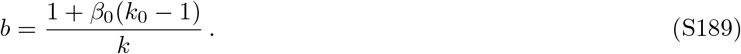

Finally, the continuity conditions at 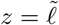 yield an analogous eigenvalue equation for 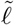:

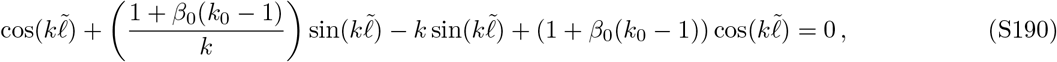

which can be written in the compact form,

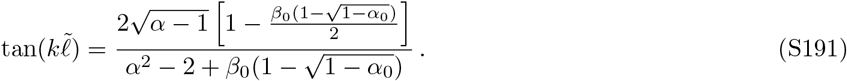

The solution to this equation,

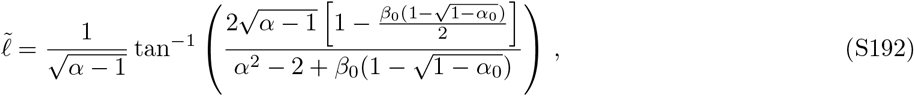

will sensitively depend on the magnitude of the compound parameter

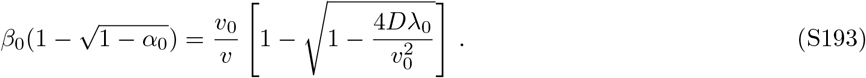

When 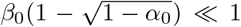, the solution for 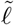 is only slightly perturbed from its value in our original model in Eq. (S142). Since we are primarily interested in cases where *v*_0_ ≿ *v*, this condition will apply whenever

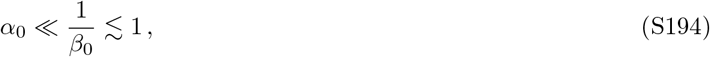

or in terms of the original parameters,

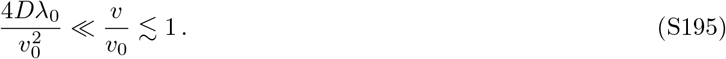

For a fixed value of λ_0_ ≾ λ, this will always be satisfied for sufficiently large *v*_0_.

On the other hand, when 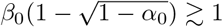, the solutions for 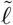 and *ρ*(*z*) will be substantially different than the original model in Supplementary Information 4.1. This can occur both when *α*_0_ is close to the washout threshold (1 — *α*_0_ ≪ 1), but also for small values of *α*_0_ provided that the flow rates are sufficiently high:

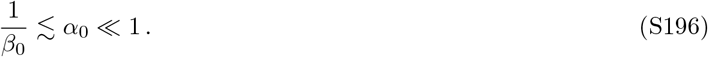

This second case corresponds to a scenario where the growth rate λ_0_ is much larger than λ, but *v*_0_ is still sufficiently high to ensure that the washout condition is still satisfied (*α*_0_ ≪ 1). This represents an inversion of the usual scenario, in which the vast majority of biomass is produced in the washed-out region due to the diffusion of cells from a stable population further downstream. While this is a potentially interesting regime, we believe that it is unlikely to apply for a typical bacterial population in the human colon. We will therefore focus on the first case 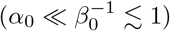 below, while leaving a full analysis of the other regimes for future work.

When 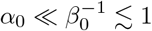, the full piece-wise solution for the density profile reduces to

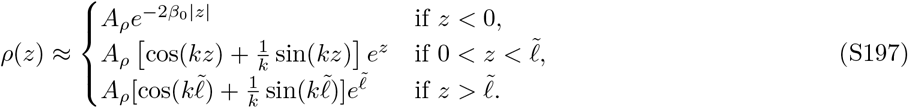

where 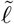 is approximately equal to the original solution in Eq. (S142). Similarly, neutral fixation probability is given by

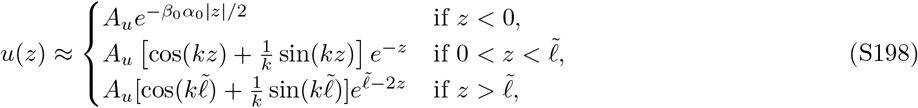

and the distribution of future common ancestors is

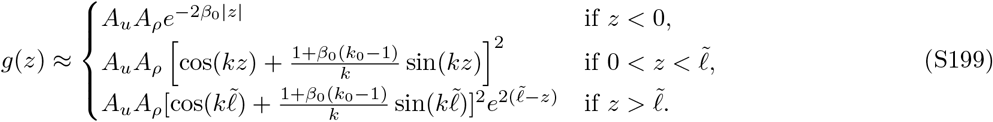

The effective evolutionary parameters can be obtained by essentially repeating the same derivation from Supplementary Information 4.2. The total population size is still dominated by the 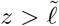 region, so the normalization constant *A_ρ_* is unchanged from before. The normalization condition, *∫ g*(*z*)*dz* = 1, now yields a slightly different expression for the normalization constant *A_u_*:

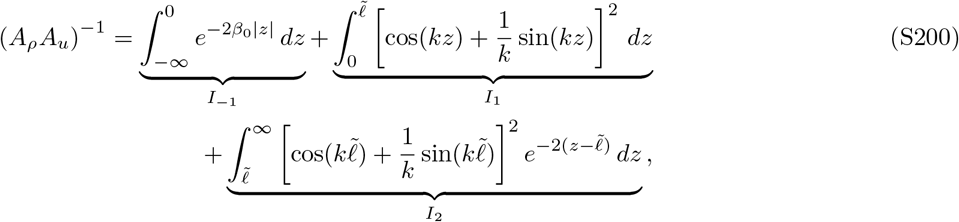

which includes an additional contribution from the integral

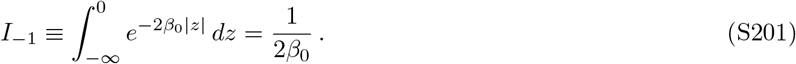

The effective growth rate can then be expressed as

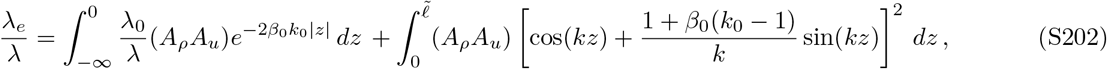

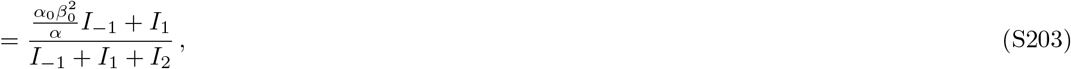

where *I*_1_ and *I*_2_ are the same as in Supplementary Information 4.2 above. Similarly, the effective population size can be written as

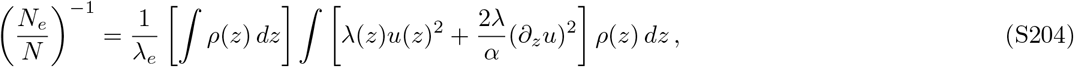

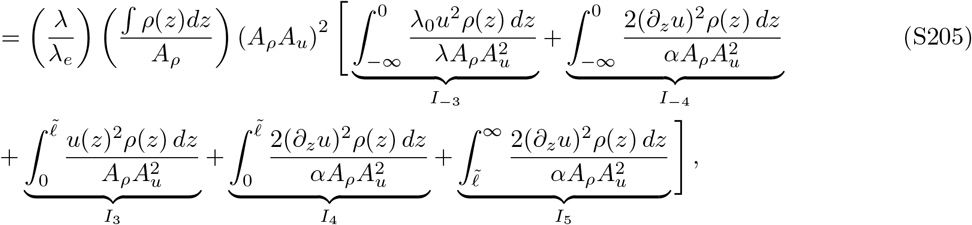

which includes two new contributions,

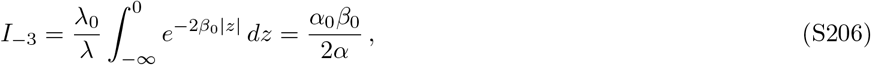

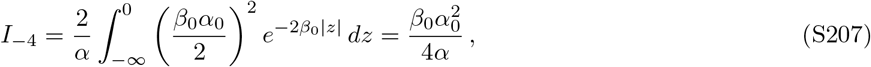

from the *z* < 0 region. Putting everything together, we find that

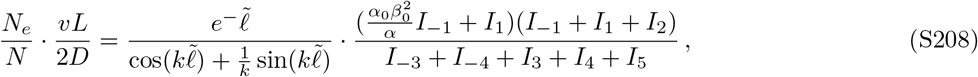

where *I*_1_, *I*_2_, *I*_3_, *I*_4_, and *I*_5_ are the same as in Supplementary Information 4.2 above. Comparing these results to our previous expressions, we see that *I*_-3_ and *I*_-4_ are smaller than *I*_3_ + *I*_4_ + *I*_5_ by an additional factor of *α*_0_*β*_0_ ≪ 1, so they make a negligible contribution to the denominator of Eq. (S208). This implies that *N_e_* is at least as large as in the original model in Fig. 2. Taken together, this shows that the addition of a strongly washed out region to an otherwise established population has only a small perturbative effect on the scaled effective population size (provided that the washed out growth rate does not become arbitrarily large). We can therefore conclude that it is the extra growth, rather than washout per se, that drives the enhanced rates of genetic drift in Fig. 2F.

#### 4. Establishment probability of strongly beneficial mutations

We calculated the establishment probabilities in Fig. 3B,C by numerically solving the differential equation in Eq. (S114) for the special case where λ_*m*_(*x*) = (1 + *s*)λ(*x*). In our minimal model, this equation reduces to the simple form,

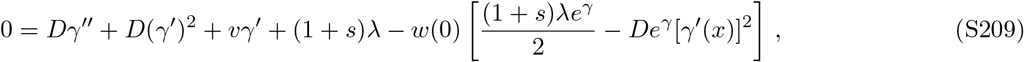

for 0 ≤ *x* ≤ *ℓ*, while the boundary conditions are given by Eqs. (S115) and (S119):

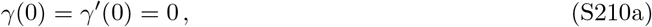

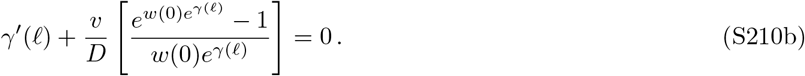

Numerical solutions to this boundary value problem were obtained using the shooting method. For a given combination of *s, λ, v*, and *D*, we scanned through a range of *w*(0) values and computed the corresponding values of *γ*(*ℓ*) and *γ*’(*ℓ*) by integrating Eq. (S209) from the initial condition *γ*(0) = *γ*’(0) = 0 using the solve_ivp function in SciPy. We then used these *γ*(*ℓ*) and *γ*’(*ℓ*) values to determine the unique choice of *w*(0) that satisfies the boundary condition in Eq. (S210b). The resulting profiles, *w*(*x*) = *w*(0)*e*^*γ*(*x*)^, were used to calculate the establishment probabilities in Fig. 3.

#### 5. Long-term growth rate of strongly beneficial mutations

The establishment probabilities in Supplementary Information 4.4 can be contrasted with the long-term growth rates of beneficial mutations derived in Supplementary Information 3.2. In our minimal model, Eq. (S122) reduces to

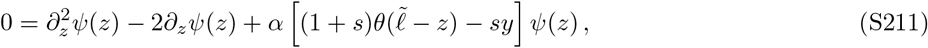

where we have adopted the same diffusion units as in Eq. (S131) above, and defined the scaled exponential growth rate

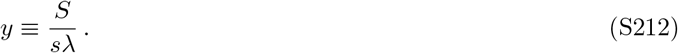

This equation admits a piecewise solution of the form,

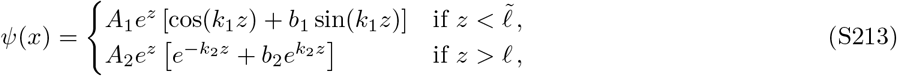

where *k*_1_ and *k*_2_ are given by

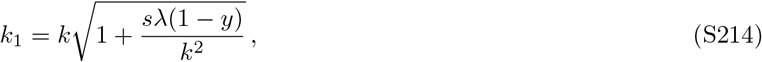

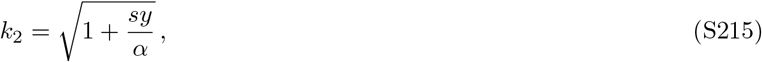

and 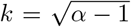 is the neutral inverse length scale defined in Eq. (S138) above. The boundary conditions at *z* = 0 and *z* = ∞ require that *b*_1_ = 1/*k*_1_ and *b*_2_ = 0. The continuity conditions at 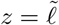 then lead to an eigenvalue equation for *y*:

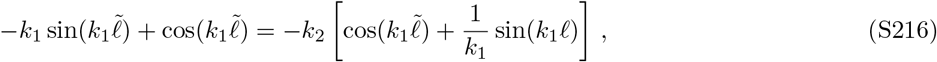

which can be rewritten in the compact form

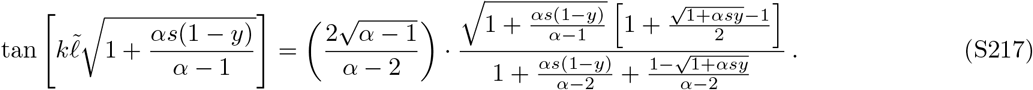

When *s* = 0, this reduces to our earlier expression in Eq. (S143) relating 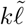 and *α*. For nonzero values of *s*, it allows us to solve for *y* as a function of *s* and *α*.

Motivated by Fig. 3, we will be particularly interested in regimes where *s* ≪ 1, but not necessarily small compared to 1/*λ_e_τ*_mix_. In this regime, we will be able to show that the *s*-dependent terms in the square roots will always be small compared to one. Expanding Eq. (S217) to first order in *s*, we obtain

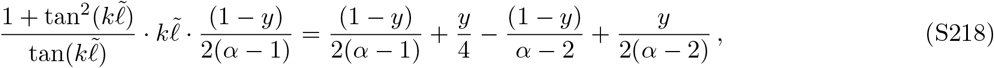

We can evaluate this expression in the diffusion-dominated growth (*α* ≫ 1) and flow-dominated growth (*α* – 1 ≪ 1) regimes respectively. In the flow-dominated regime, we have 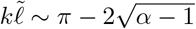, so that Eq. (S218) reduces to

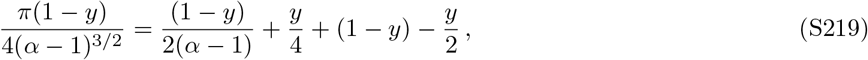

and hence

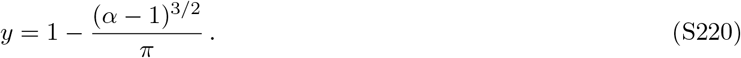

On the other hand, in the diffusion-dominated regime, we have 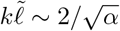 so that Eq. (S218) reduces to

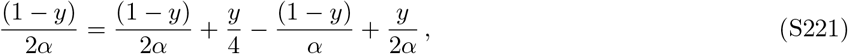

and

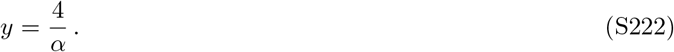

Switching back to *S*, this implies that *S* ≈ *s_e_*λ_*e*_ in both the large and small *α* limits provided that *s* ≾ 1.

### 5. APPLICATION TO BACTERIA IN THE HUMAN COLON

We sought to apply our framework to the human gut microbiota by leveraging the extensive parameter estimates in Ref. (28). The authors of this study developed a quantitative modeling framework (similar to Eq. 1) that explicitly accounts for the interplay between fluid flow, nutrient uptake, short-chain fatty acid production, and changes in local pH, while neglecting cell death [*d*(*x*) ≪ λ(*x*)]. The authors combined this computational model with *in vitro* growth measurements and published data on human physiology to obtain spatially resolved predictions for the steady-state growth rates and density profiles of two representative species in the human gut: *Bacteroides thetaiotaomicron* and *Eubacterium rectale.* We adapted these estimates for our evolutionary framework using the procedure described below.

We began by manually extracting the steady-state growth rate profiles from Fig. 3B in Ref. (28). Since these growth profiles represent solutions to a boundary value problem, we expect that small numerical artifacts in the extraction process will prevent these initial estimates from admitting a steady state solution in Eq. (2). To overcome this issue, we “refit” the extracted growth profiles by introducing a small horizontal shift

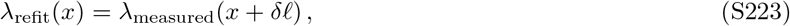

and determined the unique value of *δℓ* that yields a steady state solution for *u*(*x*) and *ρ*(*x*). We carried out this refitting step using a variant of the shooting method described in Supplementary Information 4.4, setting *s* → 0 so that *w*(*x*) becomes proportional to *u*(*x*). We verified that the fitted values of *δℓ* were small compared to the other length scales in the problem — e.g. the minimum diffusion length scale *D/v*(0), or the distance required for λ_measured_(*x*) to decay to half its maximum value — which suggests that our refitting procedure should have a negligible influence on the inferred evolutionary parameters.

Once the relevant values of *δℓ* were identified, we used the same numerical procedures to generate the steady-state solutions for *u*(*x*), *ρ*(*x*), and *w*(*x*), under the assumption that 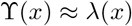. The effective evolutionary parameters in Fig. 4F-H were obtained by numerically integrating the resulting spatial profiles over the relevant values of *x*.

#### Mutation trajectory data in Fig. 5

The example mutation trajectories for the *B. massiliensis* population in Fig. 5 were obtained from our previous study (15), in which we performed deep metagenomic sequencing on 19 fecal samples from a single individual over a six month interval. Raw sequencing data was processed exactly as described in Fig. S7F of Ref. (15), producing a list of inferred mutation trajectories (Supplemental Data 1). This particular subject received a 2-week course of oral antibiotics starting on day 57, so we only plotted data from the samples that were collected prior to the start of treatment.

### 6. EXTENSIONS TO WALL GROWTH AND OTHER SPATIAL REFUGIA

Most of our analysis has focused on a simple model where all of the dynamics take place along the longitudinal axis of the lumen. In this section, we will show how our mathematical framework can be extended to account for simple forms of cross-sectional structure as well. Our use of a one-dimensional model was motivated by previous work by Cremer *et al* (26, 28), who argued that peristaltic mixing is sufficiently rapid in the human gut to destroy any cross-sectional structure within the luminal fluid itself. In particular, given an estimated effective diffusion constant of *D*≈10^6^*μm*^2^/*s*, non-motile cells will diffuse across the inner diameter of the colon (*d*~3*mm*) in ~10 minutes (26). This is significantly shorter than the rate of cell division (Fig. 4), which suggests that any cross-sectional structure will quickly average out on longer evolutionary timescales (e.g. Fig. 5).

However, this simple argument will not necessarily extend to cells in the mucosa or the intestinal crypts (25), where mixing is thought to occur more slowly. By mitigating the effects of dilution, these cells could have a natural advantage in contributing genetic material to the future population, and could therefore play an outsized role in the emergent evolutionary dynamics. While a detailed analysis of surface growth is beyond the scope of the present paper, we can gain some insight into main effects by analyzing a simpler model — designed to provide a rough approximation to wall growth in the human gut — in which a well-mixed luminal compartment is connected to *K* ≫ 1 semi-isolated subpopulations on the walls of the large intestine (Fig. S1).

We assume that each of these subpopulations (or *refugia*) contains 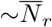. cells, which compete amongst themselves, but are otherwise isolated from the other subpopulations on the wall. These dynamics can be viewed as an approximation to the local competition that emerges from the diffusion of resources within a continuous surface population. Here we approximate these dynamics with a simple local growth rate for each refuge,

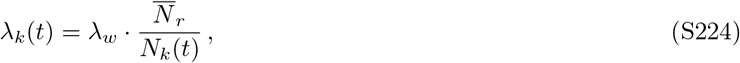

which is inversely proportional to the current number of cells, *N_k_* (*t*). We assume that the cells in the lumen compete for a separate set of resources, so that they have their own local growth rate,

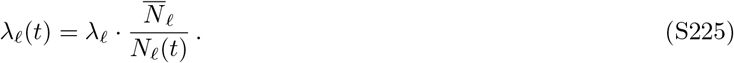

We note that the steady state growth rates λ_ℓ_ and λ_*w*_ will not necessarily be equal to each other, so that the typical generation times will vary between the lumen and the wall.

In addition to growth, we assume that the wall populations and the luminal compartment can exchange cells with each other. We will let *m*λ_*w*_ denote the per capita rate that cells migrate from the wall into the lumen, and we will let *ε*λ_*ℓ*_/*K* denote the corresponding rate of migration from the lumen into a particular subpopulation on the wall. Note that m and ε are both scaled by their local generation time, so that they can be interpreted as a traditional per generation migration probability; the parameter *ε* coincides with the *engraftment rate* defined in the main text, and will play a crucial role in our analysis below. For simplicity, we will assume that individual wall populations cannot exchange cells with each other directly, but only indirectly through diffusion into the lumen. We expect this to be a reasonable approximation for scenarios like the human gut, where active mixing makes it much easier to travel large distances within the lumen. Finally, we assume that the cells in the lumen are diluted out of the population at a per capita rate *δ*. This dilution rate is the key feature that distinguishes the lumen from the subpopulations on the walls: the latter populations only dilution indirectly by exchanging cells with the lumen. This provides them with a privileged spatial location whose effects we will investigate below.

At steady state 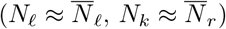, the input and output from each compartment must balance, so that

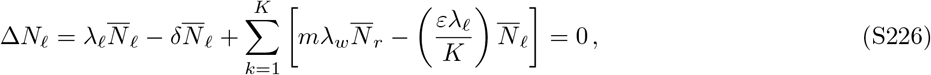

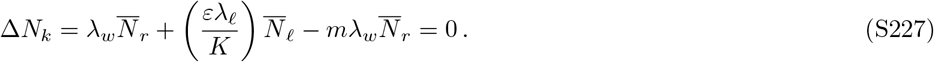

We can rearrange these equations to write them in the non-dimensional form,

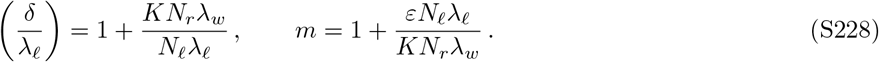

Since 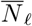 and 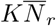 are both proportional to the longitudinal density of cells, we can also write this as

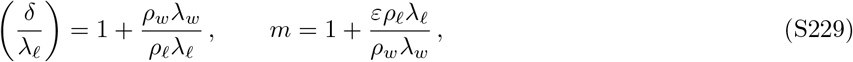

where *ρ_ℓ_* and *ρ_w_* denote the longitudinal densities in the lumen and wall respectively. Similar to our luminal models above, it will be useful to view these equations as determining *δ* and *m* as a function of the “input parameters” *ε, ρ_w_, ρ_ℓ_*, λ_*ℓ*_, and λ_*w*_. Equation (S229) shows that this relationship is mediated through two compound parameters,

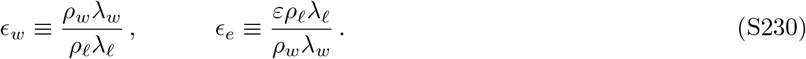

The first parameter (*ϵ_w_*) represents the ratio between the total biomass produced by wall growth and the biomass produced within the lumen. Cremer *et al* (26, 28) argued that this ratio is likely to be small for the most abundant species in the human gut, largely due to the differences in the longitudinal densities (e.g. *ρ_w_/ρ_ℓ_* ≾ 10^-3^). The ratio between λ_*w*_ and λ_*ℓ*_ is less well understood *in situ,* but it is reasonable to suppose that oxygen gradients (79, 80) and dense cellular packing in the wall (23) might limit λ_*w*_ ≾ λ_*ℓ*_ for the most abundant species that are sampled in feces (26). Based on these considerations, we will restrict our attention to regimes where *ϵ_w_* ≪ 1.

In a similar manner, the second parameter (*ϵ_e_*) can be interpreted as the ratio between the total number of cells that engraft into the wall from the lumen, and the total number produced by division within the wall itself. The empirical value of this parameter is largely unknown. Here we will focus on regimes where *ϵ_e_* ≪ 1, since this corresponds most closely to our intuitive notion of a “refuge” (i.e., more cells produced within than without). We will see below that higher engraftment rates are already effectively well-mixed with the lumen, similar to the cross-sectional structure discussed above. In terms of the underlying parameters, the assumptions that *ϵ_w_* ≪ 1 and *ϵ_e_* ≪ 1 imply that

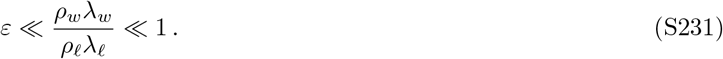

Note, however, that we make no assumptions about the *relative* values of *ϵ_w_* and *ϵ_e_*. We will see that the relationship between these two small parameters will determine the transition between refuge-dominated and lumen-dominated dynamics below.

#### Stochastic dynamics of a rare lineage

We can obtain a diffusion model for the stochastic dynamics of a rare mutant lineage by repeating our derivation in Supplementary Information 1. Let *n_k_* denote the number of mutant cells in refuge *k*, and let *n_ℓ_* the denote the corresponding number of cells in the lumen. Provided that 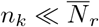 and 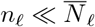, the infinitesimal change in these quantities in a timestep Δ*t* is given by

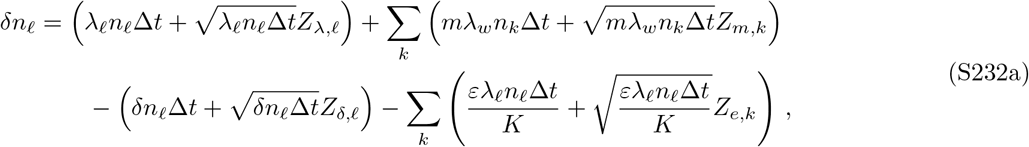

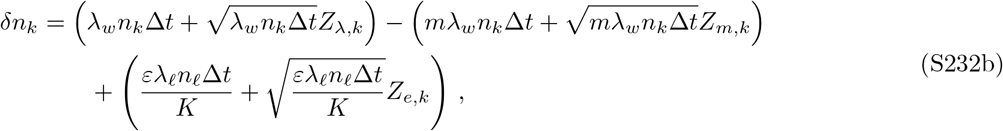

where the {*Z_i_*} again denote independent Gaussian random variables with zero mean and unit variance. After eliminating *δ* and *m* using the steady-state conditions in Eq. (S229) and enforcing the limits that *ϵ_w_, ϵ_e_* ≪ 1, these equations can be written in the compact form,

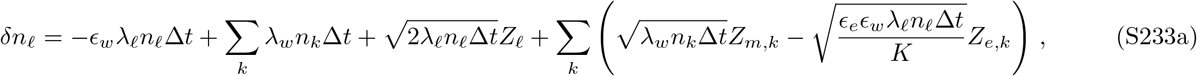

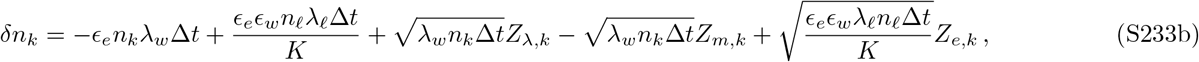

where we have merged the contributions of *Z*_λ,*ℓ*_ and *Z_δℓ_* into a single variable *Z_ℓ_*. This update rule can also be expressed in terms of the local mutation frequencies, 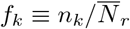 and 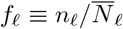, so that

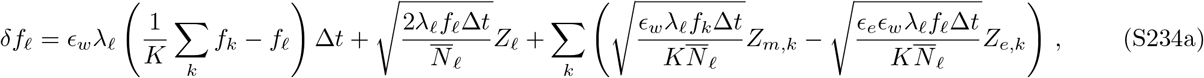

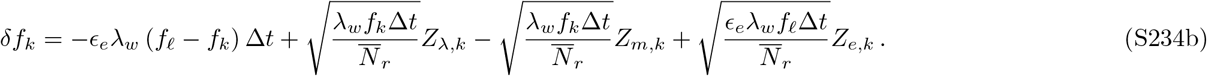

These frequency dynamics allow us to immediately identify two characteristic *mixing timescales* that equalize mutation frequencies across compartments. From Eq. (S234a), we can identify a characteristic *migration timescale,*

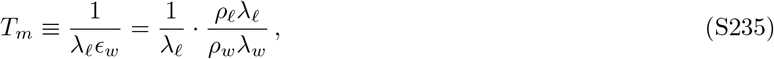

which describes how rapidly the lumen population relaxes to the average composition of the wall. Similarly, Eq. (S234b) contains a corresponding *engraftment timescale*,

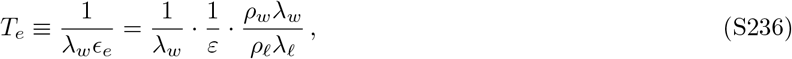

which describes how rapidly the wall populations relax to a fixed composition of the lumen. The relationship between these two timescales will play a critical role in determining the emergent evolutionary dynamics below.

Using the update rule in Eq. (S232a), we can also derive an equation for the generalization of the moment generating function,

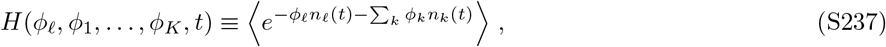

by repeating the derivation in Supplementary Information 1. This yields

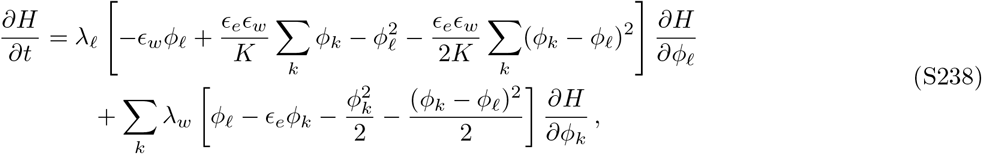

with the initial condition 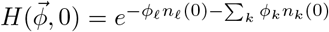.

#### Rate of genetic drift

We can use Eq. (S238) to calculate the effective rate of genetic drift by generalizing the separation-of-timescales approach in Supplementary Information 2. In this case, the linearized versions of the characteristic curves, *u_ℓ_* and *u_k_* = *u_r_*, satisfy the simple relation,

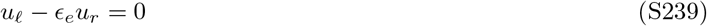

and are normalized so that 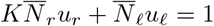. This yields

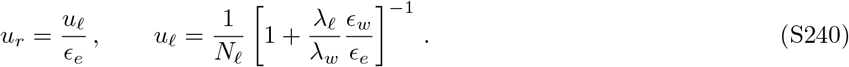

We can recognize the second term in this expression as the ratio between the mixing timescales *T_e_* and *T_m_* above. When *T_e_* ≪ *T_m_*, the value of *u_ℓ_* is completely unchanged in the presence of the wall population. In both cases, the value of *u_r_* is significantly larger than *u_ℓ_*, consistent with the notion that the refuges constitute privileged spatial locations. Our results show that the magnitude of this advantage is directly tied to the engraftment ratio *ϵ_e_*, which is consistent with our intuition above.

We can use these results to calculate the effective rate of genetic drift by repeating the perturbation expansion in Supplementary Information 2. This yields

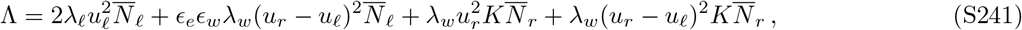

which reduces to

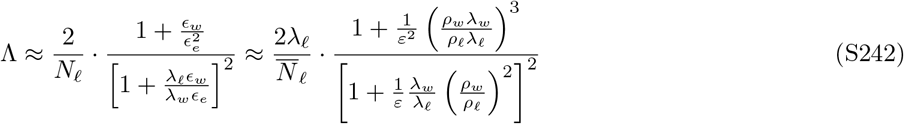

in the limit that *ϵ_w_, ϵ_e_* ≪ 1. The value of this expression critically depends on how the engraftment ratio *ϵ_e_* compares to the internal scales 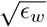 and 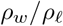, which are both small compared to one. When 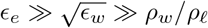, we find that

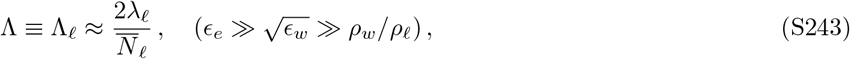

which is equivalent to the original rate of drift in the absence of the wall. In terms of the underlying parameters, this regime occurs when

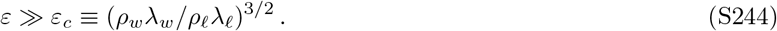

This shows that even tiny rates of engraftment are sufficient to ensure convergence to the lumen-dominated regime. In the opposite extreme, where 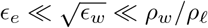, we find that

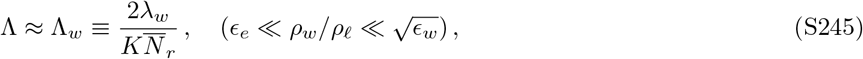

which is equivalent to the rate of genetic drift in an effective population that is the same size as the entire wall. This amplifies the rate of genetic drift by a factor of

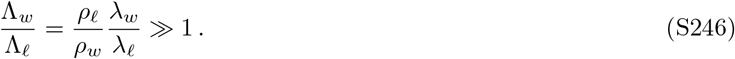

This amounts to a 1000-fold increase when *ρ_w_/ρ_ℓ_* — 10 ^3^ and λ_*w*_ – λ_*w*_. In terms of the underlying parameters, this wall-dominated regime occurs when

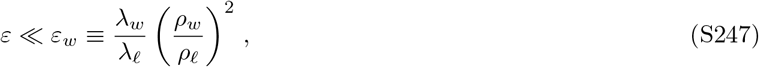

which is identical to the condition that the engraftment timescale *T_e_* in Eq. (S236) is much greater than the migration timescale *T_m_* in Eq. (S235). In other words, wall-dominated drift occurs when fluctuations in the wall propagate to the lumen more rapidly than the other way around.

In between these two extremes 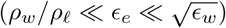, the rate of genetic drift scales as

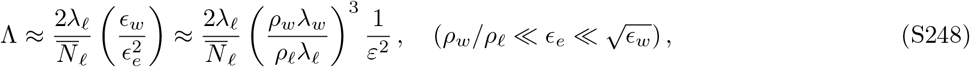

which interpolates between the Λ_*ℓ*_ and Λ_*w*_ limits above.

#### Breakdown of the separation-of-timescales approximation

The perturbation expansion used to derive Eq. (S242) will be valid on timescales much longer than the mixing timescales *T_m_* and *T_e_* in Eqs. (S235) and (S236). At a minimum, this requires that the drift timescale in the entire gut, *T*_drift_~1/Λ, must be long compared to *T_m_* and *T_e_* above. When *T_e_ T_m_*, this yields the condition,

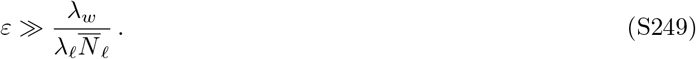

Intuitively, this condition requires that large numbers of cells must successfully engraft somewhere in the wall population during a single wall generation time. However, in addition to this global constraint, our model in Eq. (S234b) also assumed that the mutant lineage remains rare in individual refugia as well. This requires that the mixing timescale *T_e_* must also be fast compared to the drift timescale within an *individual* subpopulation, 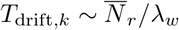. This leads to a more stringent condition,

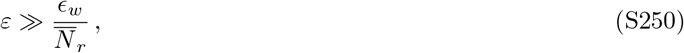

which depends on the absolute population size of an individual refuge. (Note that this is the first time where we have observed an explicit dependence on 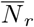, as opposed to the entire wall population, 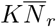.)

When 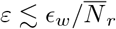, the separation-of-timescales approximation breaks down, and the dynamics are instead dominated by the slow spread of mutations across many approximately clonal refugia. We can understand these dynamics with a simple heuristic picture that is valid in the limit that 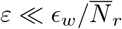. At any individual time step, each subpopulation will be fixed for either the mutant or the wildtype, and the lumen will consist of a weighted mixture of these *K* clonal populations. New lineages will enter a given subpopulation from the lumen at rate 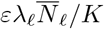. The vast majority of these lineages will go extinct; however, with probability 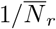, a lucky migrant will drift to fixation within the refuge on a timescale 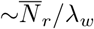 and will then propagate back into the lumen. The net effect of this event is to replace the focal subpopulation with a single individual drawn at random from one of the other *K* refuges. Thus, the dynamics in this regime will be equivalent to an effective model with *K* “individuals” that reproduce with an effective “division rate” 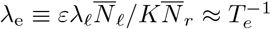. On timescales *t* ≫ *T_e_* ≫ *T_m_*, this leads to an effective rate of genetic drift,

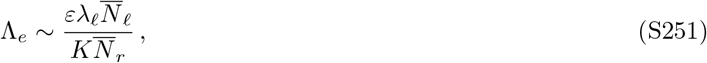

which can be substantially smaller than the wall-dominated regime above. We can now see that the origin of this slow behavior is the long time that it takes for mutations to sequentially fix across many individual refugia. This highlights an important way in which many isolated refugia are qualitatively different than a single one.

These results show that wall growth can only produce elevated rates of genetic drift within a limited parameter regime,

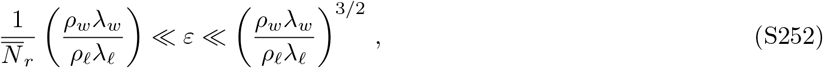

which sensitively depends on the value of 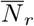. In particular, if 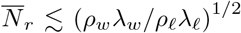, wall growth can only *reduce* the apparent rates of genetic drift. Larger rates of drift are possible above this threshold, up to a maximum possible amplification factor

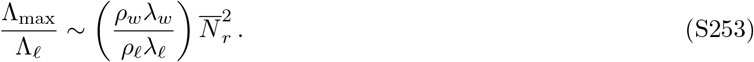

Unlike Eq. (S246) above, this bound becomes smaller for smaller values of *ρ_w_λ_w_/ρ_ℓ_*λ_*ℓ*_, and can only be counteracted by larger local refuge sizes 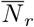. While the relevant magnitudes of 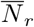 are largely unknown, these analytical results nevertheless place strong constraints on the ability of wall growth (or other spatial refugia) to enhance the rate of genetic drift. Rough estimates of the relevant parameters suggest that these effects are unlikely to explain the large shifts in frequency observed in the human fecal samples in Fig. 5. Our results also highlight the importance of migration through the lumen. This suggests that it will be impossible to fully separate the impact of wall growth from the longitudinal dynamics within the luminal fluid above. We can attain a rough correspondence between these two pictures by taking 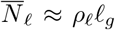, where *ℓ_g_* is the typical length scale of the ancestral region, *g*(*x*) ∝ *u*(*x*)*ρ*(*x*), in Figs 2 and 4. However, we caution that our current analysis has focused only on the simplest possible caricatures of wall growth, and omits many important features of real microbiota. A detailed understanding of these surface-growth dynamics (and their interactions with the luminal fluid above) remains an important topic for future work.

**FIG. S1.**
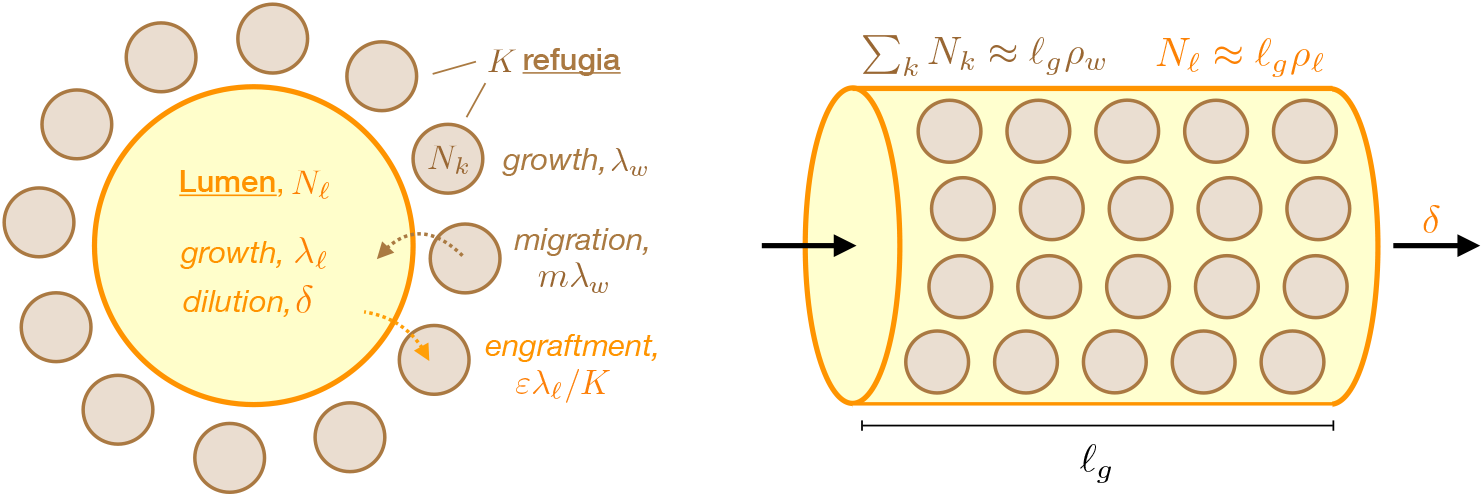
Schematic of simplified model of wall growth. A well-mixed luminal compartment exchanges cells with *K* semi-isolated subpopulations (or “refugia”) arranged on the walls. Cells move from the refugia to the lumen with a per capita migration rate *m*λ_*w*_, where λ_*w*_ is the growth rate on the wall; conversely, cells move from the lumen into the wall with a per capita engraftment rate *ϵ*λ_*ℓ*_, where λ_*ℓ*_ is the growth rate in the lumen. Cells in the lumen experience constant dilution at rate *δ,* while cells in the refugia can only be diluated if they happen to migrate into the lumen. The balance between growth, dilution, and migration leads to steady state population sizes 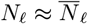 and 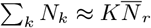 in the lumen and wall, respectively. If the luminal compartment has a cylindrical geometry with length *ℓ_g_*, these steady-state population sizes are proportional to the longitudinal densities *ρ_ℓ_* and *ρ_w_* of the lumen and wall, respectively.

